# Large-Scale Assessment of Animal-to-Human Drug Translation Using Natural Language Processing

**DOI:** 10.64898/2026.05.20.726540

**Authors:** Simona E. Doneva, Tilia R. Ellendorff, Gerold Schneider, Leonhard Held, Viktor von Wyl, T. Ian Simpson, Beate Sick, Benjamin V. Ineichen

## Abstract

**Background:** Large-scale estimates of animal-to-human drug translation and the study characteristics associated with successful translation remain limited. The expanding preclinical literature also challenges manual evidence synthesis. We developed a natural language processing (NLP) pipeline to structure and link preclinical and clinical evidence at scale.

**Methods:** In this retrospective meta-research study, we analysed more than 500,000 neuroscience-related animal drug studies from PubMed and linked them to clinical trial and regulatory approval data. NLP methods extracted drug, disease, and experimental design characteristics from abstracts and full texts. Translation was defined as progression to completed phase III/IV trials or regulatory approval. Logistic regression assessed associations between preclinical study characteristics and successful translation.

**Findings:** Among 291,624 drug entities identified in animal studies, 6·7% entered clinical development and 3·1% reached phase III/IV trials or regulatory approval. At the drug–disease level, 4·4% entered clinical development and 1·9% achieved translation. Restricting analyses to successfully linked ontology entities increased estimates to 11·3% and 4·1%, respectively. Male-only animal studies predominated, whereas reporting of randomisation, blinding, and sample size calculations remained limited. Testing across multiple species and reporting blinding were associated with higher odds of successful translation.

**Interpretation:** Only a minority of interventions tested in animals progress to advanced clinical development or regulatory approval. Greater species diversity and blinding were associated with improved translational success. NLP-based evidence synthesis may support scalable evaluation of translational research and identification of potentially modifiable research practices.

**Funding:** Swiss National Science Foundation, UZH Digital Entrepreneurship Fellowship, Universities Federation for Animal Welfare.

**Research in context:** *Evidence before this study:* We searched the literature for studies quantifying large-scale animal-to-human translation and factors associated with successful translation. Existing work was mainly limited to specific diseases, interventions, or manually curated datasets, and large-scale linkage of animal and clinical evidence remained limited.

*Added value of this study:* We developed a natural language processing pipeline linking more than 500,000 animal studies to clinical trial and regulatory approval data. The study provides large-scale estimates of translation and identifies experimental characteristics associated with successful translation.

*Implications of all the available evidence:* The findings suggest that only a minority of interventions tested in animals progress to advanced clinical development or regulatory approval. Greater species diversity and reporting of blinding were associated with improved translation. Automated evidence synthesis may support more systematic evaluation of translational research practices.

## 1 Introduction

Translating basic scientific discoveries from animals into actual patient care is a central objective and challenge of biomedical research [1, 2]. In drug development, poor translational success leads not only to substantial economic losses but also to delayed or unrealized benefits for patients. One prominent example is stroke, a major cause of disability worldwide [3]. Despite active development of drug candidates in animal models and clinical trials, effective treatment options for human diseases remain limited [4–6]. One frequently mentioned reason for low translational success is limitations in the design and conduct of preclinical research [7, 8].

These concerns center on two key aspects of scientific validity of animal research [9, 10]. One is the ability of a study to support reliable cause-and-effect conclusions. Methods such as randomization and blinding reduce bias and improve confidence in causal inference. The other is the extent to which findings apply beyond controlled laboratory conditions. This depends on whether results hold across different settings, populations, and ultimately in humans. Despite growing attention to these practices, it remains unclear if and to what extent an association between preclinical study design and translational success or failure exists.

Meta-research techniques such as evidence synthesis are an important approach for identifying gaps, patterns, and challenges in biomedical research [11, 12]. When applied to the study of animal-to-human translation, they have been used to summarize the available evidence on translational success [13, 14]. Recent efforts have also aimed to examine how characteristics of preclinical studies influence translational outcomes [15]. However, these efforts have largely been confined to narrow research domains, and the rapid expansion of the scientific literature poses challenges for large-scale evaluation of translation [16]. There is a need for approaches that enable analyses across entire biomedical fields, such as neuroscience, and ultimately benefiting translational research.

Natural language processing (NLP) techniques enable large-scale analysis of scholarly literature [17, 18]. In the preclinical domain, they have been used to map animal study characteristics such as used sexes and ages, and to extract information such as drug and disease names from study abstracts [19, 20]. However, these approaches remain fragmented and have not been integrated into a unified pipeline for systematically identifying and structuring animal evidence. In addition, they have not been extended to link the evidence bases of animal and clinical research.

The objective of this study is to quantify large-scale patterns of drug translation from animal research to clinical development. We present an automated NLP pipeline that identifies animal studies in neuroscience, extracts their experimental study design features, and links these data to clinical trial and regulatory approval records. Using this resource, we (1) analyse trends in study design characteristics in animal drug studies, (2) estimate the publicly available translation proportions of drug interventions from animal studies to clinical trials, and (3) explore associations between preclinical study design characteristics and successful translation. We release our code and the resulting curated dataset, which can be used as a starting point for further meta-research and evidence synthesis efforts.

## 2 Results

### 2.1 Included Studies

For animal drug studies, we retrieved more than 21 million PubMed articles using a neuroscience-related search string, of which approximately 6 million were eventually classified as animal studies. After applying filters for drug and disease entity detection, we identified 540,999 articles as drug-testing animal experiments. We successfully retrieved full text for 371,872 of these articles (Fig. 1 and Supplementary Fig. A1).

**Fig. 1:**
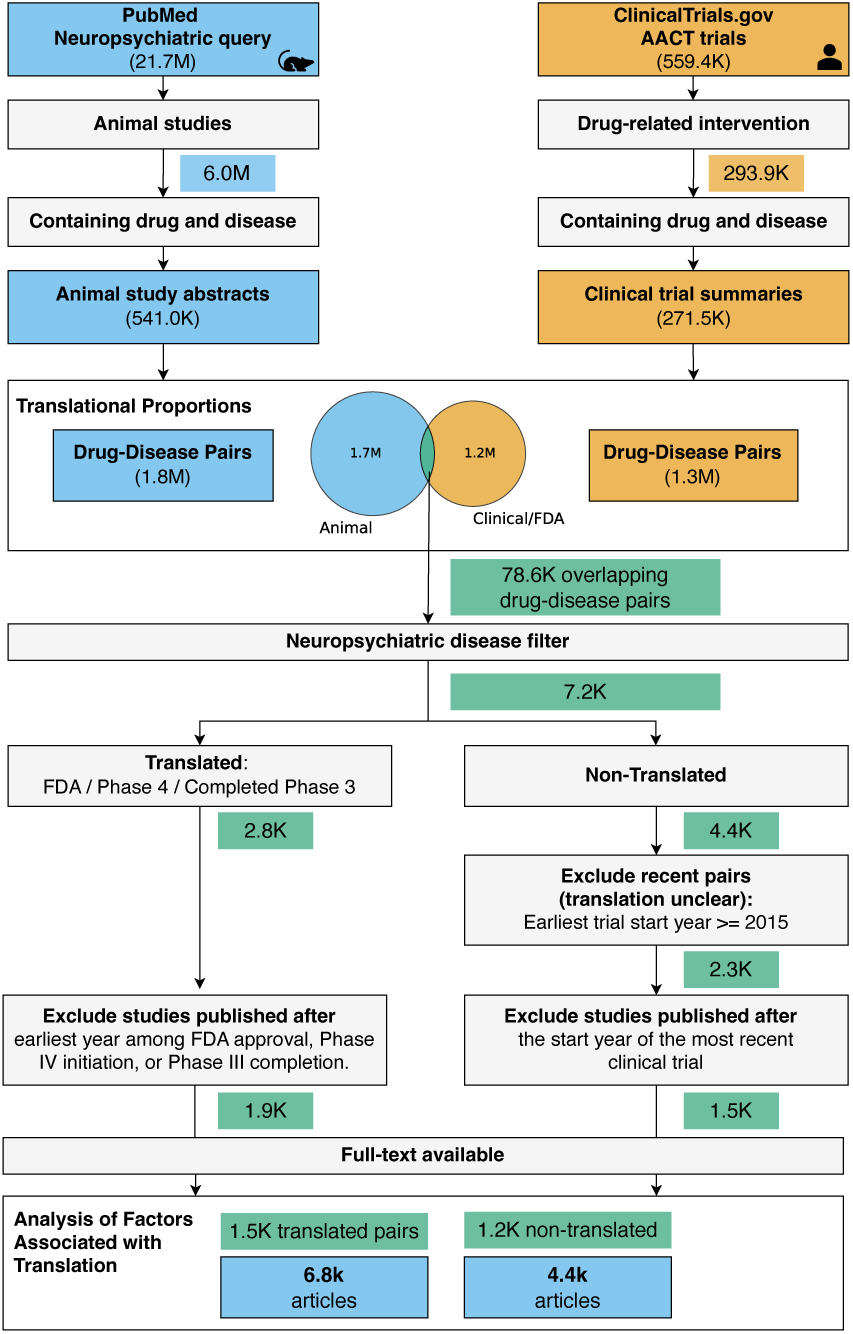
Data and analytics workflow. Translation proportions were calculated based on overlapping drug–disease pairs between clinical trials and animal studies. The drug-disease pairs were then filtering for neuropsychiatric diseases, and we selected the supporting animal studies which were published before a clinical milestone to assess factors associated with translation success.

For clinical studies, 559,371 trials were downloaded from the AACT database, of which 293,949 were interventional and involved a drug-related intervention. We identified both drug and disease names in 217,488 of these trials, which formed the core clinical dataset used in the analysis (Fig. 1 and Supplementary Fig. A1).

Additionally, we matched 2,745 FDA-approved drugs to entities identified in the animal studies. We extracted corresponding disease names from label indication information, resulting in 9,350 drug–disease pairs that we incorporated into the clinical dataset.

### 2.2 Corpus Coverage and Information Extraction Performance

To assess the coverage of our dataset, we compared our results with a reference systematic review of animal studies in multiple sclerosis. The review included 497 studies, of which 463 were indexed in PubMed. Our pipeline retained 337 of these studies (73%), with most losses attributable to unsuccessful full-text retrieval (Supplementary Table C3). Manual inspection showed that some abstracts referred only to the animal disease model (e.g., Experimental Autoimmune Encephalomyelitis) without mentioning a disease name.

BioLinkBERT was identified as the best NER model for disease, drug, and strain name recognition (Supplementary Sec. C.1). Evaluated against the PreclinIE dataset, extraction of those entities achieved F1-scores above 80%. The regular expression-based methods achieved F1-scores above 90%, with the exception of sample size detection, which showed lower recall and an F1-score of 65%. (Supplementary Table C2). Similar performance was observed when evaluating against the annotations from the published systematic review (Supplementary Sec. C.3).

### 2.3 Drugs and Diseases Entities and Normalization

For drugs, 116,529 unique mentions were extracted from the clinical corpus and 425,284 from the animal corpus (Table 1). Entity linking was successful for more than 70% of frequent mentions in both datasets, reducing them to 88,884 unique drug names in clinical trials and 291,624 in animal studies.

**Table 1:** Entity linking statistics for clinical and preclinical datasets. Linking success is shown over all entities (frequency-weighted coverage), and unique entities (distinct-entity coverage). Multiple synonyms may map to a single medical concept, resulting in overall fewer unique entities. Parent terms refers to the enriched version of the entities, where specific entities were expanded to their more generic version.

| Dataset | Type | All entities | Unique | Linking success<br>(all / unique) | After<br>linking | With parent<br>terms |
| --- | --- | --- | --- | --- | --- | --- |
| Clinical<br>(*n* = 217,488) | Drugs | 515,010 | 116,529 | 77.6%/43.1% | 88,412 | 88,884 |
|  | Diseases | 614,840 | 162,417 | 53.8%/28.6% | 122,679 | 122,807 |
| Preclinical<br>(*n* = 540,999) | Drugs | 1,499,728 | 425,284 | 70.7%/44.1% | 291,436 | 291,624 |
|  | Diseases | 1,035,001 | 123,494 | 73.9%/41.4% | 79,672 | 79,820 |

For diseases, 162,417 and 123,494 unique mentions were extracted from the clinical and preclinical corpora, respectively, and after linking reduced to 122,807 entities in clinical trials and 79,820 in animal studies. Entity linking was higher for preclinical data (73%) than for clinical data (54%). Many disease terms unique to the clinical dataset originated from Phase 1 trials and described participant status (e.g., *healthy* or *healthy volunteers*) or procedures (e.g., *cardiac surgery*, *hemodialysis*), which are not typically represented in disease ontologies and contributed to lower linking success.

### 2.4 Trends in Animal Drug Study Characteristics

There was a bias toward male animals, with 44% of studies (n = 163,617) using males only and 15% using females only (n = 55,906) (Fig. 2A). Although the absolute number of female-only studies, as well as studies including both sexes, has increased since the early 2000s, their relative proportions have remained largely stable over time. For one-third of the articles, we were unable to extract the animal sex information. Rodent models dominated the species distribution. Studies using mice only accounted for 41% of all studies (n = 151,568), followed by rat-only studies at 31% (n = 114,213) (Fig. 2B). After 2010, mouse-based studies increased rapidly, resulting in a higher proportion of studies using mice, while the number of rat studies remained relatively constant.

**Fig. 2:**
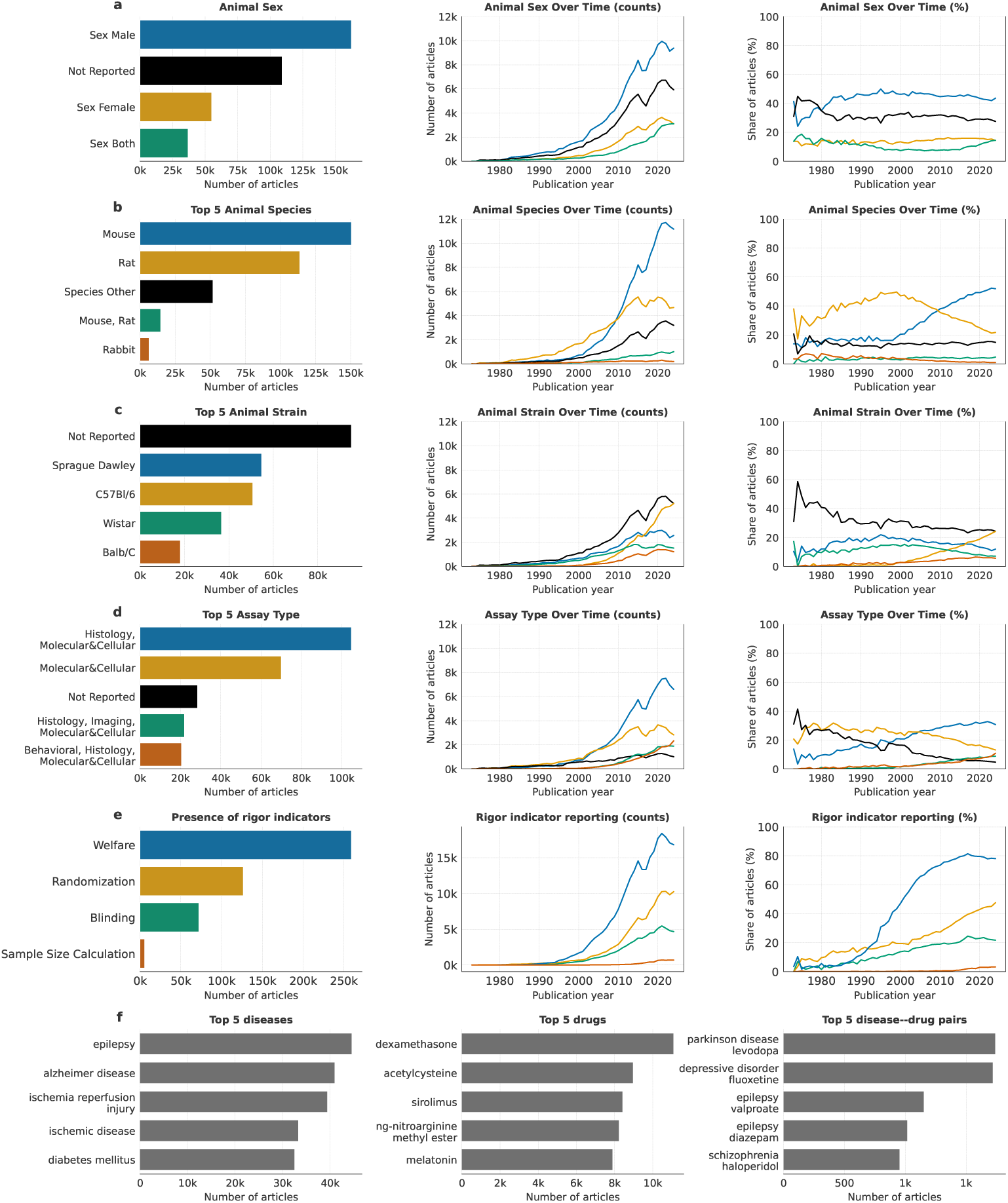
Evolution of reporting of animal characteristics and rigor indicators in the biomedical literature. For each variable, the left panel shows the total number of articles reporting the most frequent categories across the full study period. The middle panel shows the absolute number of articles per year reporting each category. The right panel shows the corresponding proportion of articles per year, normalized by the total number of articles published in that year. Only years with at least 25 articles are shown.

Outcome assessment most was most frequently reported as a combination of histological and molecular&cellular assays (Fig. 2D). Finally, the reporting of rigor indicators was overall low, with randomization, blinding, and sample size calculation being reported by 34%, 20%, and 2%, respectively (Fig. 2E).

### 2.5 Assessment of Animal-to-Human Translation

#### 2.5.1 Translational Proportions

When considering drugs independent of target disease, 19,419 of the 291,624 (6.7%) extracted from animal study abstracts also appeared in clinical records. Among these, 9,021 (3.1% of all drugs; 46.5% of those entering clinical records) were associated with a completed Phase III, any Phase IV trial and/or FDA approval. Restricting the analysis to successfully linked drug entities (n = 57,327) increased the proportion entering clinical records to 22.6%, and 11.3% that were classified as translated (Table 2).

**Table 2:** Clinical translation proportions of preclinical drug entities and drug–disease pairs. Counts are reported for all entities and only the entities successfully linked to an ontology term. Percentages indicate the fraction relative to the total preclinical count.

|  | All |  | Linked |  |
| --- | --- | --- | --- | --- |
|  | Count | % | Count | % |
| **Drugs (any indication)** |  |  |  |  |
| Total preclinical | 291,624 |  | 57,327 |  |
| Reached clinical (any phase) | 19,419 | 6.7 | 12,927 | 22.6 |
| Phase 3/Phase 4/FDA | 9,021 | 3.1 | 6,502 | 11.3 |
| **Drug–disease pairs** |  |  |  |  |
| Total preclinical | 1,807,945 |  | 648,416 |  |
| Reached clinical (any phase) | 78,638 | 4.4 | 62,425 | 9.6 |
| Phase 3/Phase 4/FDA | 33,409 | 1.9 | 26,290 | 4.1 |

When considering drugs together with their target disease, 78,638 of the 1,807,945 (4.4%; 46.5% of those entering clinical records) drug–disease pairs identified in the animal corpus appeared in clinical records, and 33,409 (1.9%) were classified as translated. After restricting to pairs with successfully linked drug and disease entities (n = 648,416), 62,425 (9.6%) appeared in clinical records and 26,290 (4.1%) reached Phase III, any Phase IV trial, and/or FDA approval (Table 2).

#### 2.5.2 Analysing Factors Associated with Translation

##### Data

From the 78,638 drug–disease pairs reaching clinical records, 7,223 remained after restricting to the list of neuropsychiatric conditions. Of these, 2,096 pairs were excluded as recent (first clinical evidence from 2015 onward). Among the remaining 5,127 pairs, 33% (1,710) had no preclinical studies published before the clinical milestone and were excluded (Fig. 3e, all studies after the orange square). A further 646 pairs had no full-text article successfully retrieved for any associated study (Table F12). The final dataset comprised 1,565 translated and 1,206 non-translated pairs (Fig. 1 and Fig. 3c,d).

**Fig. 3:**
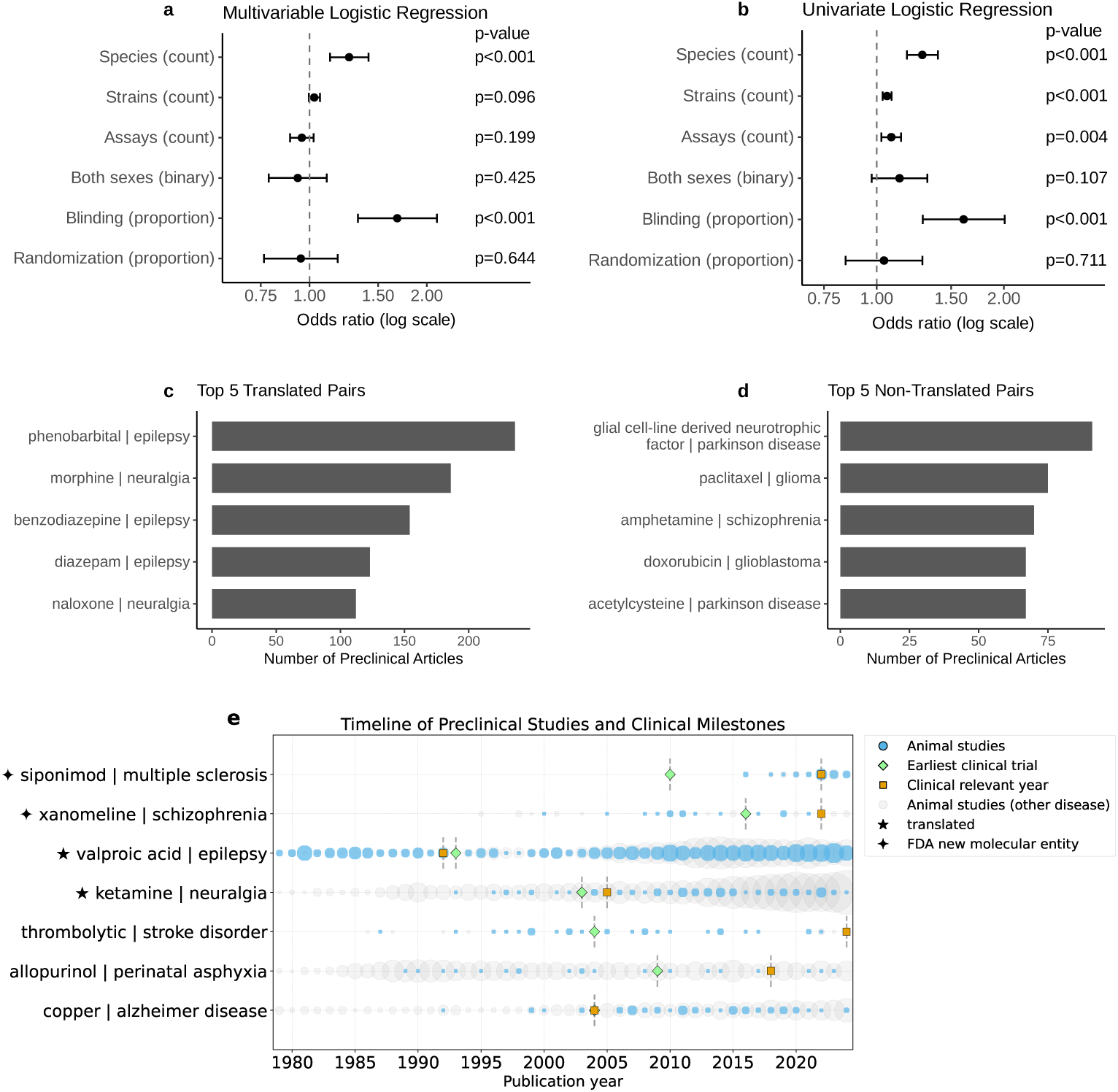
Associations between preclinical study characteristics and translation success. Adjusted odds ratios from multivariable (a) and univariate (b) logistic regression models; bars indicate 95% confidence intervals. Panels (c) and (d) show the top translated and non-translated drug–disease pairs, ranked by the number of supporting preclinical studies. Panel (e) presents timelines of preclinical studies and clinical milestones for five randomly selected drug–disease pairs, including two recently approved FDA new molecular entities. Bubble size reflects the number of publications per year.

##### Descriptive Statistics

Testing a drug in more than two countries occurred in 55.8% of translated drug–disease pairs compared with 45.9% of non-translated pairs. Similar patterns were observed for testing in more than two species (50.5% vs 42.5%) and more than two strains (56.2% vs 49.0%). In addition, at least one study reporting blinding was present in 54.6% of translated pairs compared with 42.7% of non-translated pairs (Table 3).

**Table 3:** Comparison of translated and non-translated drug–disease pairs across preclinical study design diversity and reporting rigor. All design and rigor metrics are reported as number and percentage of drug–disease pairs. The Δ column reports percentage-point differences (translated minus non-translated).

| Characteristic | Approved | | Non-approved | | $\Delta$<br>pp |
| --- | --- | --- | --- | --- | --- |
|  | Count | % | Count | % |  |
| Number of drug–disease pairs | 1,565 |  | 1,206 |  |  |
| Unique drugs | 708 |  | 698 |  |  |
| Unique diseases | 107 |  | 98 |  |  |
| Total number of studies | 6,839 |  | 4,359 |  |  |
| Tested in both sexes | 715 | 45.7 | 514 | 42.6 | 3.1 |
| Tested in $\geq 2$ species | 790 | 50.5 | 512 | 42.5 | 8.0 |
| Tested in $\geq 2$ strains | 880 | 56.2 | 591 | 49.0 | 7.2 |
| Tested with $\geq 2$ outcomes | 1,283 | 82.0 | 967 | 80.2 | 1.8 |
| Tested in $\geq 2$ countries | 873 | 55.8 | 553 | 45.9 | 9.9 |
| Blinding reported ($\geq 1$ study per pair) | 854 | 54.6 | 515 | 42.7 | 11.9 |
| Randomization reported ($\geq 1$ study per pair) | 931 | 59.5 | 649 | 53.8 | 5.7 |
| Welfare reported ($\geq 1$ study per pair) | 1,307 | 83.5 | 997 | 82.7 | 0.8 |

##### Analysis of Associations using Logistic Regression

In univariate logistic regression, testing drugs in a greater number of species and strains, a larger number of assays, and having a higher proportion of studies reporting blinding were associated with higher odds of successful translation (Fig. 3a). In multivariable logistic regression, testing in a greater number of species and a higher proportion of studies reporting blinding remained associated with successful translation (OR = 1.26, 95% CI: 1.13–1.42, *p <* 0.001; OR = 1.68, 95% CI: 1.33–2.12, *p <* 0.001; Fig. 3b). Supplementary Sec. F.3 provides results from additional sensitivity analyses.

## 3 Discussion

In this study, we quantified large-scale patterns of drug translation from animal to clinical development using an automated NLP pipeline linking animal studies to clinical trials and FDA approvals. We found that only a minority of drug interventions reported in animal studies are represented in subsequent clinical trials or regulatory approvals. Our analyses further revealed trends in study design characteristics across animal research and identified experimental features associated with a higher likelihood of translation from preclinical studies to clinical testing, including testing a drug in more than one species and the use of blinding procedures.

The observed male sex bias is consistent with long-standing patterns in the biomedical literature. In neuroscience, male-only studies have been reported to outnumber female-only studies by approximately 6:1 [21], closely matching our findings. Although we observe a modest increase in studies including both sexes, consistent with prior work [22], the gap is closing only slowly. This raises concerns about the generalizability of findings to women, given the continued underrepresentation of female animals in drug testing [23–25].

We also observe a marked increase using mice as experimental model species over the past two decades, in line with previous analyses [26, 27]. This trend, contrasted with the relatively stable use of rats, likely reflects early advances in gene-targeting and genomic technologies in mice [26]. However, rats may provide more translationally relevant models for complex behaviors and pathophysiology [28]. More broadly, the strong reliance on rodent models may limit biological diversity and translational relevance [26, 29].

We observed modest improvements over time in the reporting of randomization, blinding, and sample size calculations [30, 31]. These findings are consistent with previous studies that suggest that many animal studies continue to face challenges in implementing and reporting rigorous experimental designs [32–34].

Based on the constructed dataset, we estimate a translation proportion of 3–11% for drugs independent of the target disease and 2–4% for drugs considering their target disease (drug–disease pairs). These estimates lie between the 0% reported for mental health disorders and the 7% reported for nervous system diseases in a related study assessing translation of both drug and non-drug therapeutic interventions [14].

Our findings further suggest that several experimental design characteristics in animal research are associated with translational success. In particular, testing in multiple species and the use of blinding were associated with higher odds of translation.

Evaluating interventions across multiple species may improve the generalizability of preclinical findings by reducing the likelihood that observed effects are specific to a single animal model [26, 35]. Similarly, the use of blinding reduces the risk of observer bias and may lead to more reliable estimates of treatment effects [36, 37].

Supporting the relevance of methodological quality, a recent analysis across three disease areas reported a higher prevalence of statistical misapplication in animal studies preceding unsuccessful clinical trials [15]. Together with our findings, this suggests that aspects of preclinical experimental design and conduct can be quantified and studied in relation to translational outcomes.

Nonetheless, the results presented here should be interpreted in light of some important limitations. First, these associations do not imply causation. In observational research, “reverse causality” remains a possibility: it is plausible that candidates with high early potential attract more funding, which in turn allows for more extensive multi-species testing and higher methodological rigor [38]. To minimize these effects, we used a strict temporal filter, analyzing only the preclinical evidence that was available before clinical milestones were reached, which excluded a substantial proportion of included studies.

Second, translation was defined based on the progression of drug–disease pairs to late-stage clinical trials or regulatory approval. Alternative definitions, such as concordance between animal and human outcomes, may yield different estimates [13]. Furthermore, as drug–disease pairs were identified through co-mentions in abstracts, they may not capture direct experimental relationships or distinguish multiple experiments within a study. Future work could address these limitations by incorporating more advanced structured evidence extraction techniques [39, 40].

Third, translational processes are complex and rarely follow a linear pathway. Animal studies may occur alongside or after early human research, and publication can be delayed. Prior work shows that many animal studies are published only after clinical testing has begun or even after approval [4, 41, 42]. Moreover, some interventions, such as drug repurposing, may proceed without new animal studies and clinical development may be discontinued for reasons unrelated to efficacy, including commercial considerations, or strategic reprioritization.

Fourth, although this is, to our knowledge, the largest analysis of its kind, its scope is constrained by the data sources used. For preclinical research, other literature resources such as Embase and emerging preclinical study registries could contain relevant studies [43, 44]. For clinical evidence, publications reporting trial results could be integrated alongside registry data, and additional registries beyond ClinicalTrials.gov could improve global coverage. Restricted access to full-text articles for automated retrieval is another limitation. In addition, as neuroscience is a large and heterogeneous domain, the generalizability of these findings to other fields remains to be established. Finally, concerns regarding research integrity, including image or results manipulation, and paper mill activity, have been increasingly reported in the biomedical literature [45–47]. In addition, this study was also likely limited by publication bias. Studies with positive or significant findings are more likely to be published than negative or null results in both the animal and clinical domain [48, 49]. The impact of these issues is difficult to assess and represents a broader challenge for evidence synthesis.

Despite these limitations, our results highlight the potential of NLP for structuring and linking evidence across animal and human drug development. The proposed computational pipeline can substantially reduce the effort required to analyze large volumes of animal research literature, enabling the identification of trends and gaps in the evidence base, and allowing to study its links to clinical drug development. It may also support indication-specific subgroup analyses, facilitating the investigation of whether certain animal models translate more effectively to humans. Furthermore, as demonstrated by our analysis of associations between study design characteristics and translational outcomes, this approach can help identify potentially modifiable research practices to improve translation. While the precise relationship between experimental design choices in animal research and downstream clinical outcomes needs further investigation, we hope that our work will motivate future research in this area.

## 4 Methods

### 4.1 Study Design

Retrospective observational meta-research study using NLP to extract structured information from full-text animal drug development studies and link these data to corresponding clinical trial registry and regulatory drug approval records.

### 4.2 Data Sources and Study Selection

#### 4.2.1 Animal Studies

We sourced animal studies from PubMed and applied three successive filters to identify (1) studies in the field of neuroscience, (2) studies involving animals, and (3) studies investigating a drug intervention for a specific disease.

To identify studies related to neuroscience research, we applied a broad search strategy, retrieving records up to December 18, 2024.^1^ To identify animal studies, we used a recently published dataset of annotated PubMed titles and abstracts to train a binary classifier for study type (animal vs. other) [50]. Finally, drug and disease mentions were extracted from the titles and abstracts of the articles. Only studies containing both entities were retained for further analysis.

Full texts were initially retrieved from the PubMed Central (PMC) Open Access subset, a freely accessible archive of biomedical articles [51]. For articles not included in this subset, we used Cadmus, an automated full-text retrieval system that accesses publisher content from Elsevier and Wiley through institutional license keys [52]. The “Methods” section of each article was then extracted, and split into individual sentences to facilitate traceability of the extracted information.

#### 4.2.2 Clinical Trials and FDA Data

Clinical trial information was obtained from the Aggregate Analysis of ClinicalTrials.gov (AACT) database [53].^2^ We first filtered the database to include clinical trials involving drug-related interventions based on the study metadata. We then extracted drug and disease mentions from the study official titles and brief summaries, as described in Section 4.3.2. We retained only records in which we detected both a drug and a disease entity, thereby restricting the dataset to studies that test drug interventions for specific conditions (see Supplementary Sec. A for details).

To improve identification of successful clinical translation, we incorporated drug information from regulatory datasets from the U.S. Food and Drug Administration (FDA)^3^ (further details in the Supplementary Sec. D).

### 4.3 Information Extraction

#### 4.3.1 Animal Studies Data Extraction

We extracted animal study characteristics from PubMed article’s abstracts, full texts, and metadata. From PubMed metadata, we obtained publication year, journal name, and first author’s institutional affiliation. We mapped author affiliations to geographic locations using a recently published geoinference method [23].

From the abstracts, we extracted drug and disease names using a fine-tuned named entity recogniton (NER) model on the PreClinIE dataset [54]. For subsequent analysis, we aggregated entity mentions at the document level and retained only unique entities. From the Methods sections of full texts, we obtained information on animal species, sex, randomization, masking, animal welfare, sample size calculation, and outcome measures. For the extraction of those elements from the text, we extended a recently published regular expression–based (regex) library [55] (see Supplementary Sec. B for assay library construction). Animal strain and total animal counts were extracted from the Methods section with a NER model.

#### 4.3.2 Clinical and FDA Data Extraction

Building on our prior work, we used the language model BioLinkBERT [56] to extract drug and disease names, achieving F1-scores of 0.90 for drug entities and 0.85 for disease entities in an independent test set [57]. The clinical trial’s phase, status, and start, completion, and first submission year were extracted directly from the AACT database.

To extract drug–disease relationships from FDA data, we integrated information from drug approval records and associated drug labels. For each drug, we compiled all associated indications (diseases) together with their respective years of approval, accounting for the fact that a single drug may have multiple approvals for distinct indications. Further details are provided in Supplementary Sec. D.

### 4.4 Drug and Disease Entity Linking

The linkage between animal and clinical/FDA data was based on shared drug and disease names. A major challenge in this step is the substantial variability in biomedical terminology. For example, “stroke” and “cerebrovascular accident”, or “cariprazine hydrochloride” and “cariprazine hcl”, being the same entities. To standardize these variations, we applied named entity linking to map NER-extracted text spans to normalized identifiers in biomedical ontologies [58]. For drug entities, our target ontology was the Metathesaurus from the Unified Medical Language System (UMLS) [59]. For disease entities, we used the Monarch Disease Ontology (MONDO) [60]. Extracted mentions were linked to ontology concepts using SapBERT^4^ [61].

Another challenge concerns the granularity of entity mentions. Specific terms such as “relapsing-remitting multiple sclerosis” or “177Lu-DOTA-rituximab” captured fine-grained subdomain distinctions but complicated aggregation for high-level trend analysis. Therefore, we retrieved their parent concepts from the ontology hierarchy. In this example “multiple sclerosis” and “rituximab” would be added as entities associated with the article. Further details on both entity linking and parent aggregation are provided in Supplementary Data E.

### 4.5 Performance Evaluation

#### 4.5.1 Information Extraction

We evaluated the performance of the information extraction modules using precision, recall, and F1-score. For the information identified using regular expressions, performance was evaluated directly on the full PreClinIE dataset as no fine-tuning of the libraries was necessary [54].

The best-performing NER model was identified using 10-fold cross-validation on the PreClinIE dataset, comparing BioLinkBERT, BioBERT, and BiomedBERT [56, 62, 63]. Final performance is reported as the mean across all folds for the selected model. For drug and disease mentions, a prediction was considered correct if the model identified a relevant entity at least once within an abstract, rather than requiring correct identification of every individual mention. Partial matches for drug and disease entities were also counted as correct.

#### 4.5.2 Validation with a Published Systematic Review

To evaluate the reliability of our system in identifying relevant studies, we compared its outputs with a recent systematic review of drug interventions for multiple sclerosis and associated animal study characteristics [41]. We used the manually extracted animal study data provided by the authors via the study’s public repository.

Two aspects were evaluated. First, we assessed the coverage of our dataset by measuring the overlap between articles included in this review and those identified by our pipeline. Second, we evaluated annotation accuracy by comparing model-predicted study characteristics with the manually extracted annotations reported in the review.

### 4.6 Data Analysis

#### 4.6.1 Trends in Animal Study Characteristics

We reported the prevalence of animal study characteristics and indicators of methodological rigor for the PubMed articles with available full texts. We presented the most frequently observed characteristics and show trends over time using the raw number of articles, as well as the proportions of articles exhibiting specific characteristics.

#### 4.6.2 Animal-to-Human Translation

##### Definition of Translation

We tracked how far drug therapies identified in animal studies progressed along the translational spectrum. Translation was considered successful if a therapy was associated with at least one completed Phase III or Phase IV clinical trial, or with regulatory approval by the FDA.

##### Estimation of Translational Proportions

The aim of this analysis was to estimate the proportion of entities found both in animal studies and clinical trials, and the proportion of successful translations. This analysis was based on the drug and disease entities extracted from the animal study abstracts, and we used the complete animal and clinical datasets.

At the drug level, the results reflect the proportion of drugs identified in animal studies that enter clinical trials or receive approval for any indication. At the drug–disease level, this reflects whether a drug investigated for a specific indication enters clinical trials or achieves approval for that same indication.

##### Factors Associated with Translation

The aim of this analysis was to use logistic regression to identify experimental design features from animal studies associated with successful vs unsuccessful animal-to-human translation. Analyses were conducted at the drug–disease pair level and included the animal studies for which a full-text was available (additional details and experiments: Supplementary F)

###### Data

We applied several filters to obtain the final data used for the analysis (Fig. 1). First, at the drug–disease pair level, we retained only pairs where the disease matched a list of neurological and psychiatric diseases^5^. This restriction was applied because the animal studies corpus was constructed around neuroscience-related research, and we aimed to ensure that the majority of relevant animal evidence was represented in the dataset. From the remaining drug–disease pairs, those meeting the above criteria were classified as translated. Then, drug–disease pairs with recently initiated clinical activity were excluded, defined as those with a first clinical trial start year in or after 2015, since translational outcomes could not yet be determined given typical drug development timelines. The remaining pairs were classified as not translated.

Second, animal studies published after the relevant clinical milestone were excluded. This restriction was applied in order to focus only on the body of animal research that was available for translational decision making. For translated pairs, we excluded animal studies published after the earliest year of FDA approval, Phase IV initiation, or Phase III completion. For non-translated pairs, we excluded animal studies published after the initiation of the most recent clinical trial. Drug–disease pairs were excluded if the associated preclinical corpus did not contain any eligible animal studies after this filtering step.

###### Independent and dependent variables

The dependent variable captured translational success at the drug–disease pair level and was coded as 1 if the pair was supported by at least one completed Phase III or Phase IV clinical trial or FDA approval, and 0 otherwise.

Following prior work, independent variables were defined to capture key features of the preclinical evidence base related to study rigor and the range of experimental conditions [41, 64]. Variables were computed at the drug–disease pair level by aggregating information across all associated animal studies.

To characterize the range of experimental conditions, we considered four measures: the number of unique animal species tested, the number of unique animal strains used, the number of distinct assay types, and whether both sexes were represented. A pair was coded as including both sexes if male and female animals were present in the supporting evidence base, either within a single study or across multiple studies.

To capture features related to study rigor and potential bias, we assessed the reporting prevalence of two practices: randomization and blinding. For each drug–disease pair, these variables were calculated as the proportion of associated studies in which keywords related to random allocation or blinded outcome assessment were detected.

###### Estimation procedures

We first summarized the distribution of experimental characteristics across drug–disease pairs using descriptive statistics. These summaries were compared between translated and non-translated pairs. Logistic regression models were used to evaluate associations between experimental factors and translation status. Each predictor was first assessed using univariate logistic regression, followed by a multivariable logistic regression including all predictors simultaneously. Associations were reported as odds ratios (ORs) and adjusted odds ratios (AORs) with 95% confidence intervals (CIs). The logistic regression analyses were performed in R (version 4.3.3).

## Data availability

The datasets analyzed during the current study are available from:

- https://zenodo.org/records/19354961 (animal studies corpus and translation data)
- https://zenodo.org/records/19351891 (ClinicalTrials.gov)

Additional data are available from the corresponding author upon reasonable request.

## Code availability

Code for data processing, analysis, and visualization is publicly available at:

- https://github.com/Ineichen-Group/Preclinical_DrugDisease_Translation_Pipeline (animal studies processing and translation analysis)
- https://github.com/Ineichen-Group/ClinicalTrials_DrugDisease_Pipeline (Clinical-Trials.gov processing)
- https://github.com/Ineichen-Group/FDA_DrugDisease_Pipeline (FDA approved drug and indications processing)

## Supplementary information

Supplementary information is available as appendix for this paper and includes additional figures, tables, and methodological details supporting the findings of this study.

## Acknowledgements

We thank the CAMARADES research group, Peter Kind, David Price, and Douglas Armstrong (University of Edinburgh), and Samuel Pawel (University of Zurich) for helpful discussions. We also thank Michael Yates for support in data acquisition.

## Declarations

### Ethics approval and consent to participate

Not applicable. This study is based on analysis of publicly available data and does not involve human participants or new animal experiments.

### Consent for publication

Not applicable.

### Materials availability

No new materials were generated in this study.

### Author contributions

Conceptualization: B.V.I., S.E.D., and T.I.S. Methodology: all authors. Implementation: S.E.D. Validation: B.V.I. and B.S. Data curation: S.E.D. Visualization: S.E.D. Writing - original draft: S.E.D. Writing - review and editing: all authors. Project administration: S.E.D. and B.V.I. Funding acquisition: B.V.I. All authors reviewed and approved the final version of the manuscript.

## Appendix A Included Studies

Overview of the filtering steps of the clinical and preclinical datasets is provided in Fig A1.

### A.1 Details on included clinical trials

From the AACT ClinicalTrials.gov database, we selected studies with drug-related interventions. Study metadata, including the ClinicalTrials.gov identifier (nct id), brief title, official title, start date, completion date, first submission date, study phase, and recruitment status, were extracted from the studies table and linked to study descriptions from the brief summaries table.

Intervention information was obtained from the interventions table, retaining only intervention types DRUG, DIETARY SUPPLEMENT, BIOLOGICAL, COMBINATION PRODUCT, GENETIC, and OTHER. Intervention names and intervention types were aggregated at the study level. Condition names were extracted from the conditions table and aggregated at the study level.

### A.2 Metadata Trends

Metadata on the preclinical studies, including journal distribution and first author countries, are shown in Figure A2. The dataset is distributed across a large number of journals. Although a few journals such as *PLoS One* (9,965 articles) and *Scientific Reports* (7,245 articles) contribute the highest absolute number of publications, their relative share remains small, as shown in panel (a). Publication activity in the top journals increases over time, particularly after 2010, but their proportion of the total output remains limited.

Clear geographic trends are observed in panel (b). China (90,014 articles) and the USA (82,367 articles) account for the largest share of publications, with China showing a substantial increase in output in recent years. In contrast, countries such as Japan, Germany, and the UK contribute smaller but relatively stable shares over time. The proportion of articles from China increases steadily, while the relative contribution from the USA declines, indicating a shift in the geographic distribution of preclinical research output.

## Appendix B Assay Library Construction

Annotations describing outcome measures used to assess intervention efficacy showed low inter-annotator agreement in PreClinIE and were therefore unsuitable for training a prediction model. To address this limitation, we developed an independent library of commonly used outcome assessment techniques and incorporated it into a rule-based extraction approach.

The outcome domains used to characterize intervention efficacy are summarized in Table B1. These domains were defined as part of a harmonized assay vocabulary used in our classification pipeline. The core set of assays was curated through manual review of the literature, using the articles listed in Table B1 to identify commonly used readouts across domains.

Extracted assay mentions were normalized into canonical names and assigned to one of five outcome domains. To improve coverage of lexical variation, canonical assays were expanded with synonymous expressions generated using a large language model (ChatGPT) and subsequently reviewed. The resulting vocabulary was used to construct a domain-specific synonym dictionary for automated information extraction, where matches were identified using regular expressions and mapped to canonical assay names and their corresponding domains.

The outcome domains capture distinct levels of biological organization and measurement modality:

- **Behavioral**: Overt animal actions or responses, including motor, sensorimotor, cognitive, social, and affective or pain-related behaviors (e.g., rotarod, open field, elevated plus maze).
- **Imaging**: *In vivo* medical imaging techniques such as MRI, PET, and CT. This category excludes microscopy and photographic images.
- **Histology**: Tissue-based analyses, including staining methods (e.g., LFB, Nissl, H&E) and microscopy-based assessments.
- **Physiology**: Measurements of vital signs, metabolic and functional readouts, body composition, pharmacokinetics, and survival (e.g., heart rate, blood pressure, EEG, EMG).
- **Molecular & Cellular**: Assays that quantify or characterize molecular components (DNA, RNA, proteins, metabolites) or cellular properties, including gene expression and protein-based assays such as qPCR, RNA-seq, ELISA, mass spectrometry, and flow cytometry.

## Appendix C Preclinical Pipeline Evaluation

### C.1 NER Model Selection

Model performance was evaluated using 10-fold cross-validation on the PreClinIE dataset [54]. Each record in PreClinIE corresponds to either the abstract or the methods section of a PubMed article. Folds were generated with GroupKFold using a document-level grouping key derived from the document identifier, ensuring that sections from the same publication were assigned to the same fold. Each split consisted of 1,296 training samples and 144 test samples. Drug and disease entities were modeled jointly within a single NER model, while strain entities were modeled in a separate task. Models were fine-tuned using the *Hugging Face Transformers* library, and evaluation metrics (precision, recall, and F1) were computed on the held-out fold and logged during training. Final results are reported as mean scores across all folds. BioLinkBERT achieved the highest overall performance across both annotation tasks (Fig. C3). In the drug–disease setting, it outperformed BioBERT and BiomedBERT across all three metrics. In the strain setting, BioLinkBERT again had the highest recall and F1. On this basis, BioLinkBERT was selected for downstream analyses. BioBERT was not benchmarked on the strain task because it had already shown the lowest performance in the drug–disease evaluation.

### C.2 Performance on PreClinIE for All Tasks

The performance of the information extraction pipeline was evaluated using the PreclinIE dataset. Table C2 provides an overview of the extraction approaches used for each information item together with their precision, recall, and F1-scores. For drug and disease entities, we report partial-match scores. Partial matches were identified using approximate string matching implemented with difflib.get_close_matches, where a predicted entity was counted as correct if it matched any target entity with a similarity cutoff of 0.6.

### C.3 Performance Evaluated on a Published Systematic Review

We extracted the set of PMIDs from a published systematic review of animal studies of approved and failed MS disease-modifying therapies [41]. As summarized in Table C3, we tracked retention of this reference set across all pipeline stages using successive set intersections. Losses were minimal during query retrieval, animal filtering, and NER-based drug–disease identification, while the largest reduction occurred at the full-text stage, resulting in 72.8% of the original reference set remaining.

For the animal characteristics extraction performance evaluation, we used the annotations provided in the HERMES-MS repository^6^. Our evaluation dataset included 337 studies. Model performance for sex and species extraction is shown in Figure C4. The confusion matrices demonstrate strong agreement between true and predicted labels for both animal sex (Fig. C4A) and species (Fig. C4B), with most observations concentrated along the diagonal. Note that in this dataset, female animals are more frequently represented. This reflects a common experimental practice in multiple sclerosis (MS) research, where female animals are more susceptible to developing Experimental Autoimmune Encephalomyelitis (EAE), the most widely used mouse model of MS.

Manual review of five randomly sampled label–prediction discrepancies for the sex variable indicates that most mismatches arose from incomplete or ambiguous reporting of animal sex in the methods section (Table C4). Notably, one case reflected a ground-truth annotation inconsistency rather than a model misclassification, suggesting that the observed discrepancies do not exclusively represent model errors.

For the risk-of-bias elements, we compared our predictions against the annotations provided in a separate file of the HERMES-MS repository^7^. A total of 64 studies with corresponding annotations were also present in our dataset and were included in this validation analysis. Based on the confusion matrices shown in Figure C5, overall model performance appears strong for welfare, blinding, and sample size. In the matrix for “randomization” (panel B) shows a higher number of false negatives (11 cases labeled as “present” in the ground truth but predicted as “not-reported”), indicating reduced sensitivity for this category.

However, manual inspection (Table C5) indicates that several apparent false negatives are likely correct model predictions. Many positive labels were from references to human studies or general trial procedures rather than explicit random allocation of animals. Our extraction criteria require animal-related keywords in the local textual neighborhood, which likely explains part of the discrepancy and suggests that the target annotation for randomization may have been comparatively permissive.

## Appendix D FDA Data Acquisition and Processing

### D.1 Data Sources

FDA regulatory data were obtained from the openFDA platform on 11 November 2025. We downloaded (i) the Drugs@FDA application files (https://open.fda.gov/apis/drug/drugsfda/download/) and (ii) structured drug label files (https://open.fda.gov/apis/drug/label/download/). The Drugs@FDA dataset provides application-level regulatory and submission metadata, while the Drug Label dataset contains structured labeling text, including approved indications and clinical study descriptions.

### D.2 Application-level Preprocessing

All JSON files were parsed and flattened into tabular format. From Drugs@FDA, we retained application number, submission status, approval year, sponsor name, submission class, and active ingredient information, and additional drug related fields: brand name, generic name, substance name. Analyses were restricted to applications having an approved FDA status.

### D.3 Drug Name Normalization

Drug names were standardized by combining active ingredient, substance name, and brand name fields. The dataset was expanded such that each row represents a single drug–application pair. Drug strings were normalized to Unified Medical Language System (UMLS) concepts using a neural named entity normalization model that returned standardized terms and Concept Unique Identifiers (CUIs).

### D.4 Extraction of Indication Text

From the Drug Label files, we extracted application numbers and the full indications and usage section. Records without valid application numbers were discarded. Indication text was concatenated into a single string per label. We retained both the full indication text and the first indication sentence, which serves as a standardized summary of approved use.

Application-level drug records from the previous step were merged with this cleaned label data using application number as the linking key.

To derive structured disease entities from free-text indication statements, we used a large language model (LLM) using the vLLM framework for faster inference. We downloaded and applied the DeepSeek-R1-Distill-Qwen-32B model with a maximum context length of 8,192 tokens and fixed sampling parameters to ensure consistent outputs. For each drug–application record, the cleaned indication sentence was provided as input, and the model was prompted to extract the primary disease or condition entities referenced in the approved indication. When multiple indications were returned, they were separated into individual rows. Extracted indication strings were subsequently mapped to MONDO using the same normalization framework applied to drug entities.

To validate the LLM’s performance, we manually annotated 50 randomly sampled indication texts with the target disease names, yielding 201 total target disease entities (190 unique). Model outputs were compared against these annotations using both exact and relaxed matching. As shown in Table D6, we report entity counts alongside precision, recall, and F1 scores. The relaxed matching is based on normalized substring overlap, allowing matches when one entity appears within another, thereby accounting for minor variations in phrasing.

### D.5 Illustrative FDA Application-level Records

Table D7 illustrates representative drug–application records after merging with cleaned label text and applying LLM-based indication extraction. The column label_indications_first_sent contains the standardized first indication sentence extracted via rule-based parsing. The column LLM extractions shows structured disease entities returned by the LLM. This example demonstrates how the same active ingredient (cladribine) can receive approval for distinct clinical indications across different application years. Such variation is important when analyzing approval timing, as the approval year is specific to a drug–indication pair rather than the molecular entity alone.

## Appendix E Entities Linking

### E.1 Reference Ontologies

#### E.1.1 MONDO Disease Ontology

##### Included vocabulary

MONDO was selected as the primary disease ontology because it integrates disease concepts from a wide range of biomedical vocabularies [60]. The main Web Ontology Language (OWL) vocabulary was obtained in April 2025 from https://mondo.monarchinitiative.org/pages/download/. The Python library *pronto* (https://pronto.readthedocs.io/en/stable/) was used to process the ontology and extract disease terms and their synonyms. A total of 30,158 unique terms were identified (term IDs starting with MONDO:). The synonyms for each term were collected, excluding synonyms with scope “BROAD”. This resulted in a vocabulary of 128,591 unique disease names and their variations.

**Example:** Synonyms for autoimmune hepatitis (MONDO:0016264), and their scope in brackets:

- autoimmune hepatitis (EXACT)
- AIH (RELATED)
- chronic autoimmune hepatitis (NARROW)
- autoimmune chronic active hepatitis (NARROW)
- autoimmune hepatitis with centrilobular necrosis (NARROW)
- autoimmune liver disease (BROAD) -*>* **excluded**
- autoimmune chronic hepatitis (RELATED)

##### Postprocessing

MONDO can have repetitive disease entities, and the need for consolidating related disease terms has been discussed in earlier work [99]. Following the approach introduced in that prior work, we grouped disease concepts with highly similar names into unified nodes using string-matching rules. This method identifies diseases that share a common root name but differ only by stage, grade, subtype, or minor linguistic variation, and merges them into a single representative concept. For example, terms such as Alzheimer disease 2, Alzheimer disease 3, and Alzheimer disease type 1 were all grouped under the common root Alzheimer disease.

#### E.1.2 UMLS and DrugBank for Drug Mapping

##### Included vocabulary

The Unified Medical Language System (UMLS) provides a repository of biomedical and health-related concepts. We retrieved the “2024AB Full UMLS Release Files” on 28 October 2024 in ‘.RRF’ format, , which can be obtained from https://www.nlm.nih.gov/research/umls/licensedcontent/umlsarchives04.html.

Several filters were applied to the database. An overview is given in Table E8. The unique drug code names remaining after filtering numbered 474,316. Because the UMLS assigns multiple terms to each unique identifier, capturing name variants originating from different source vocabularies, this expanded to an additional 560,190 terms.

Although UMLS includes the DrugBank standard name and its synonyms, several DrugBank fields are not represented, including *International Brand Names*, *Product Names*, and *External IDs*^8^. We therefore obtained the DrugBank XML release on 15 November 2025 under an academic license from https://go.drugbank.com/releases/latest. We processed the file and extracted these fields. Then we merged the corresponding synonyms with the UMLS terms that contained a matching DrugBank identifier. This contributed an additional 93,270 terms.

The *External IDs* field was not present in the XML base file. External IDs represent “codes and identifiers used by other resources and companies, often used before choosing a marketing name.” Instead, we retrieved them directly from the DrugBank website, which yielded 15,181 additional terms. For example, the values “HSP-130” and “MYL-1401H” would be mapped to the drug “Pegfilgrastim”.

#### E.1.3 UMLS Relations Mappings

We leveraged additional UMLS relationships to further refine mappings to canonical forms. Specifically, we processed the relations provided in *MRREL.RRF* and selected two relevant relationship types: *has tradename* and *has lab number*. For example, the drug Integrilin (C0950902) is linked via *has tradename* to its canonical name Eptifibatide (C0253563). In such cases, when an entity is initially matched to Integrilin, it is finally mapped to the canonical form Eptifibatide.

### E.2 Linking with SapBERT

#### E.2.1 Embeddings

We used the Self-alignment Pretraining for BERT (SapBERT) model from the Huggingface library, pre-trained on PubMedBERT full texts, without further fine-tuning or change to the hyperparameters^9^.

We generated SapBERT embeddings for all concept terms and synonyms across the target ontologies. For each named entity, we computed its corresponding SapBERT vector and retrieved the most similar ontology concepts by ranking candidates according to their Euclidean distance (cdist) [100].

#### E.2.2 Linking Similarity Threshold Estimation

The “cdist” value can be interpreted as an indicator of the match’s probability, with larger distances suggesting lower confidence in the match. We sought to identify an optimal cdist threshold above which entities should not be linked to the ontologies, as such matches are likely to be false positives. To determine this cutoff, we manually annotated a random sample of 100 drug entities and 100 disease entities, and evaluated how different threshold values affected their classification.

The highest F1 scores were obtained at a cdist threshold of 9.65 for disease, achieving an F1 score of 0.82 (**Figure E6 (b)**). For drug entities, the highest F1 score of 0.81 was achieved at a cdist threshold of 8.20 (**Figure E6 (a)**).

At lower cdist thresholds, the model was more stringent, accepting only very close matches. This resulted in higher precision but lower recall, as the model missed some true matches that had a higher Euclidean distance. Conversely, at higher cdist thresholds, the model was less strict, which increased recall by including more true matches, but also decreased precision due to the inclusion of more false positives.

### E.3 Term to Parent Mapping

Our NER system outputs fine-granular annotations, often identifying highly specific drug and disease entities. For downstream aggregation, however, it is necessary to map these specific variants back to a common canonical or root form.

Within UMLS, we rely on the relationships provided in the MRREL table to perform this normalization. Specifically, we use the narrower (RN) relationship type (REL) and further filter by the relationship labels (RELA) supplied by the source vocabularies, which specify the precise semantic nature of each parent–child link. Only RELA values relevant to drug and chemical concepts were retained. The selected relation types and illustrative examples are shown in Table E9.

To assign dataset-specific parent concepts, we first collect all UMLS terms that appear in the dataset and treat this set as the pool of potential parents. For each detected entity, we then traverse the UMLS relation graph to retrieve all candidate parent CUIs. A parent is only accepted if it is itself present in the dataset, ensuring that mappings reflect relationships within the observed data rather than the full ontology. To prevent overly generic abstractions, we exclude parents with more than 20 children (e.g., broad categories such as Antiviral Agents).

### E.4 Entity Linking Examples

To illustrate how heterogeneous NER strings are linked to a common representation, Table E10 gives examples from the preclinical dataset. Table E11 summarizes the most frequent mapped and not-linked drug and disease entities, highlighting both successful normalization and common entities for which no ontology concept met the similarity threshold.

### E.5 Entity Overlaps Animal–Clinical Data

The overlap of unique drugs, diseases, and drug–disease pairs between the animal and clinical study corpora is shown in Figure E7.

For drugs, a total of 88,884 unique drug names were extracted from clinical trials, compared to 291,624 from animal studies (Table 1). Examples of drug entities that appear in the animal dataset but not in the clinical dataset include *MK-801*, *SB 203580*, and *PD-98059*. Entity linking for drug mentions was successful for more than 70% of frequent mentions in both datasets.

For diseases, 122,807 unique names were extracted from clinical trials and 79,820 from animal studies. Closer inspection revealed that many disease entities unique to the clinical dataset originate from Phase 1 trials and describe participant status rather than pathological conditions, such as *healthy* or *healthy volunteers*. Additionally, procedure-related terms such as *cardiac surgery* and *hemodialysis* appeared only in the clinical dataset. These entities are typically not linked to disease ontologies, which contributed to the lower entity-linking success observed for disease terms in the clinical domain (54%).

## Appendix F Factors Associated with Translation

To examine the association between preclinical study characteristics and subsequent clinical translation, we performed logistic regression analyses at the drug–disease pair level. The unit of analysis was the individual drug–disease pair.

### F.1 Data Preparation

#### F.1.1 Temporal Filtering

To avoid temporal leakage, preclinical evidence was restricted to studies published prior to the relevant clinical milestone for each drug–disease pair. This temporal filtering ensured that all preclinical predictors used in the regression models reflected information available prior to advanced clinical development.

For translated pairs, only preclinical articles published on or before the first relevant clinical milestone (i.e., the earliest Phase 3/Phase 4 trial or regulatory approval year) were retained. For non-translated pairs, preclinical articles were restricted to those published on or before the most recent recorded clinical trial start year.

#### F.1.2 Data Transformation

Preclinical studies were initially represented at the study level. Table F13 illustrates two example records corresponding to the same drug–disease pair. These records were then aggregated at the drug–disease pair level by combining information across all supporting studies.

### F.2 Variables

#### F.2.1 Dependent Variable

The dependent variable was *translation status*, defined as a binary indicator of whether a drug–disease pair achieved advanced clinical development. Pairs were classified as translated (1) if supported by at least one completed Phase 3, or Phase 4 clinical trial, or regulatory FDA approval, and as non-translated (0) otherwise.

#### F.2.2 Independent Variables

Independent variables were grouped into measures of external validity, internal validity, and evidence volume. All independent variables were computed at the drug–disease pair level by aggregating information across all supporting preclinical studies associated with each pair.

**External validity** was operationalized using four measures reflecting the diversity of the preclinical evidence base: (i) the number of unique animal species tested, (ii) the number of unique animal strains tested, (iii) the number of unique assay types employed, and (iv) an indicator for inclusion of both sexes. Note that a pair was coded as testing both sexes if the supporting evidence base included at least one study using male animals and at least one study using female animals (not necessarily within the same study).

**Evidence volume** was included as a control variable to account for differences in the number of supporting preclinical articles across drug–disease pairs. Pairs with more studies have a higher probability of exhibiting greater diversity or reporting of rigor practices purely due to increased sampling. Evidence volume was therefore measured as the total number of unique supporting articles (PMIDs) per pair and was log-transformed in the regression model.

**Internal validity** was assessed using reporting intensity of key risk-of-bias practices. Specifically, we calculated the proportion of supporting articles reporting blinding and the proportion reporting randomization for each drug–disease pair.

All models report odds ratios with 95% confidence intervals.

### F.3 Additional Results

#### F.3.1 Detailed Regression Results

Table F15 provides the numerical results underlying the forest plots in the main paper (Fig. 3a,b). In univariate analyses, a greater number of species, strains, and assays tested, as well as higher reporting of blinding, were associated with increased odds of translation. In multivariable models, the number of species tested and reporting of blinding remained significant predictors, whereas other variables were attenuated and no longer statistically significant.

As reporting practices in preclinical research have evolved over time, we next examined whether the publication year of included animal studies differed between translated and non-translated drug–disease pairs.

As shown in Fig. F8, non-translated studies were published slightly more recently (median 2010, IQR 2004–2015; *n* = 4359) compared to translated studies (median 2008, IQR 2001–2013; *n* = 6839). The distributions show substantial overlap but indicate a modest temporal shift between groups.

#### F.3.2 Adjusting for Publication Volume

Drug–disease pairs with more preclinical publications may have a higher experimental diversity because they have been studied more extensively. To assess whether the associations observed in the main analysis were driven by differences in publication volume, we performed an additional analysis incorporating the number of supporting preclinical articles per drug–disease pair as a covariate.

For each pair, we quantified the total number of supporting preclinical articles and included the logarithm of this count (log(articles)) in the multivariable logistic regression model together with the previously defined measures of external validity (species diversity, strain diversity, assay

Table F16 summarizes the distribution of supporting articles for translated and non-translated drug–disease pairs. Figure F9 shows the distribution of evidence volume across translation status (a) and the adjusted regression estimates when publication volume is included in the model (b).

Including publication volume in the regression model had only a modest impact on the estimated associations. The number of supporting articles was positively associated with translation success (AOR = 1.14; 95% CI 0.98–1.32; *p* = 0.081), although this effect was not statistically significant. Overall, the inclusion of this variable did not alter the direction of the associations observed for the other predictors.

#### F.3.3 All Drug–Disease Pairs

The preclinical evidence dataset used in this study is primarily focused on neurological disorders. However, when constructing the set of drug–disease pairs, some pairs correspond to conditions outside this domain. For such pairs, much of the relevant pre-clinical evidence may lie outside the curated dataset, potentially leading to incomplete extraction of supporting studies. Therefore, the main analysis was restricted to a subset of neurological and psychiatric conditions. To assess how including all drug–disease pairs regardless of biomedical domain affects the results, we repeated the analysis using the full set of pairs.

Figure F10 presents the regression results for this analysis. In the full dataset, several predictors were significantly associated with translation success. Species diversity and blinding remained positively associated with translation, whereas assay diversity, inclusion of both sexes, and randomization were associated with lower odds of translation.

---

**Fig. A1:**
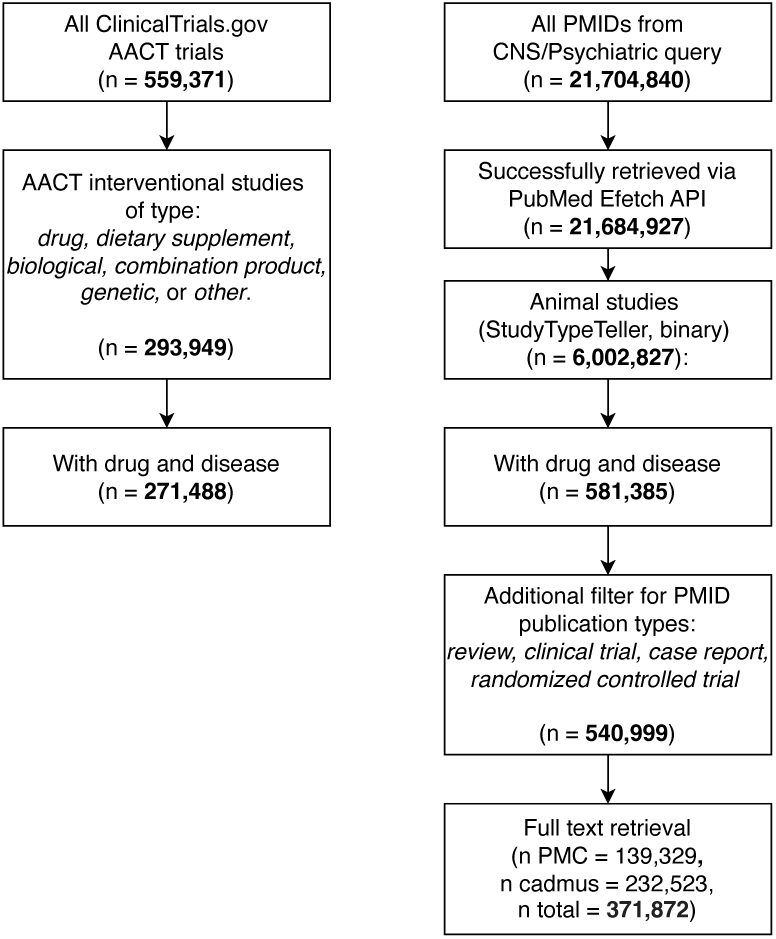
Overview of the process of relevant studies identification.

**Fig. A2:**
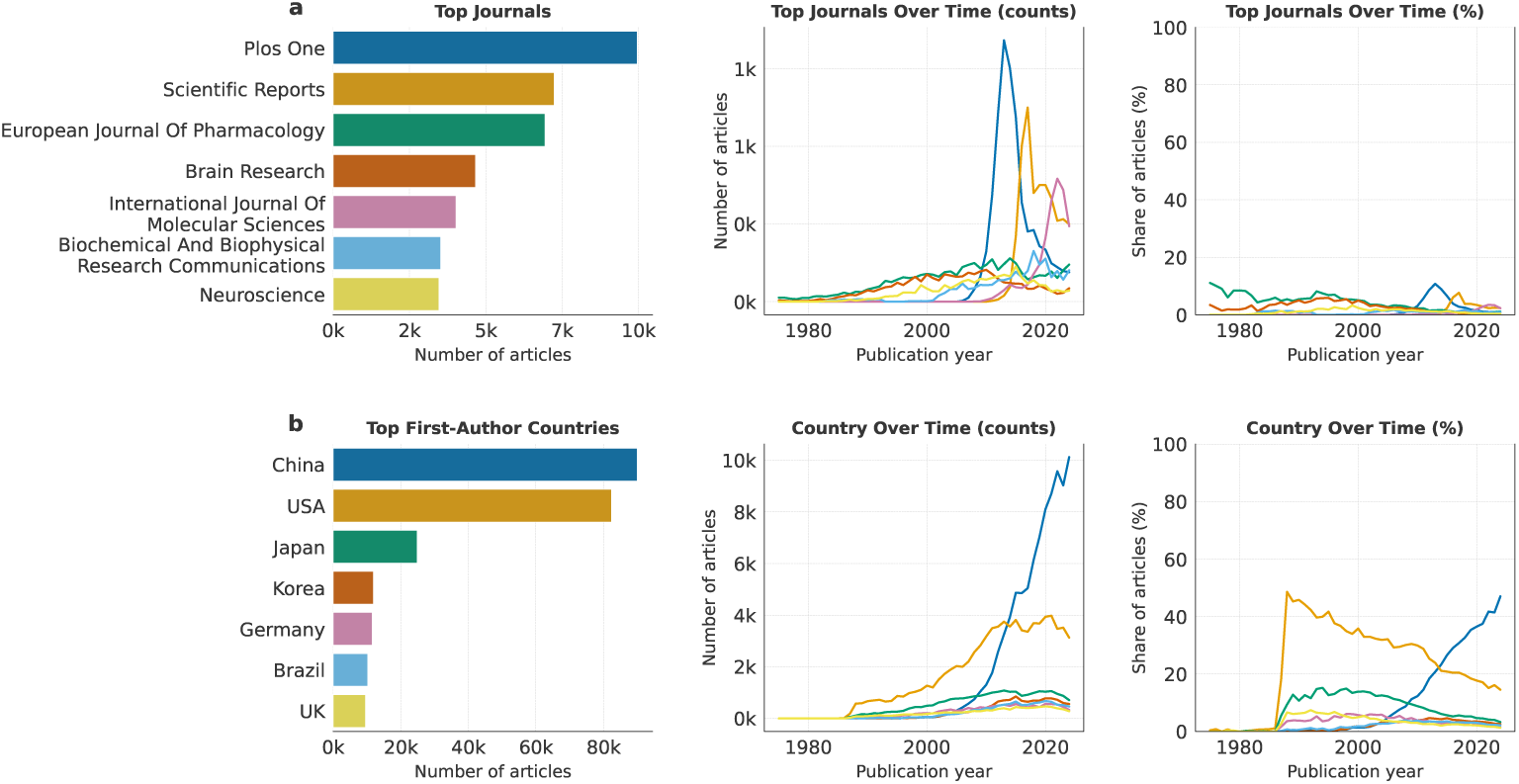
Distribution and temporal trends of journals and first author countries in the preclinical dataset. For each variable, the left panel shows the total number of articles for the most frequent categories across the full study period. The middle panel shows the number of articles per year, and the right panel shows the corresponding proportion of articles per year, normalized by the total number of articles in that year. Only years with at least 100 articles are included.

**Fig. C3:**
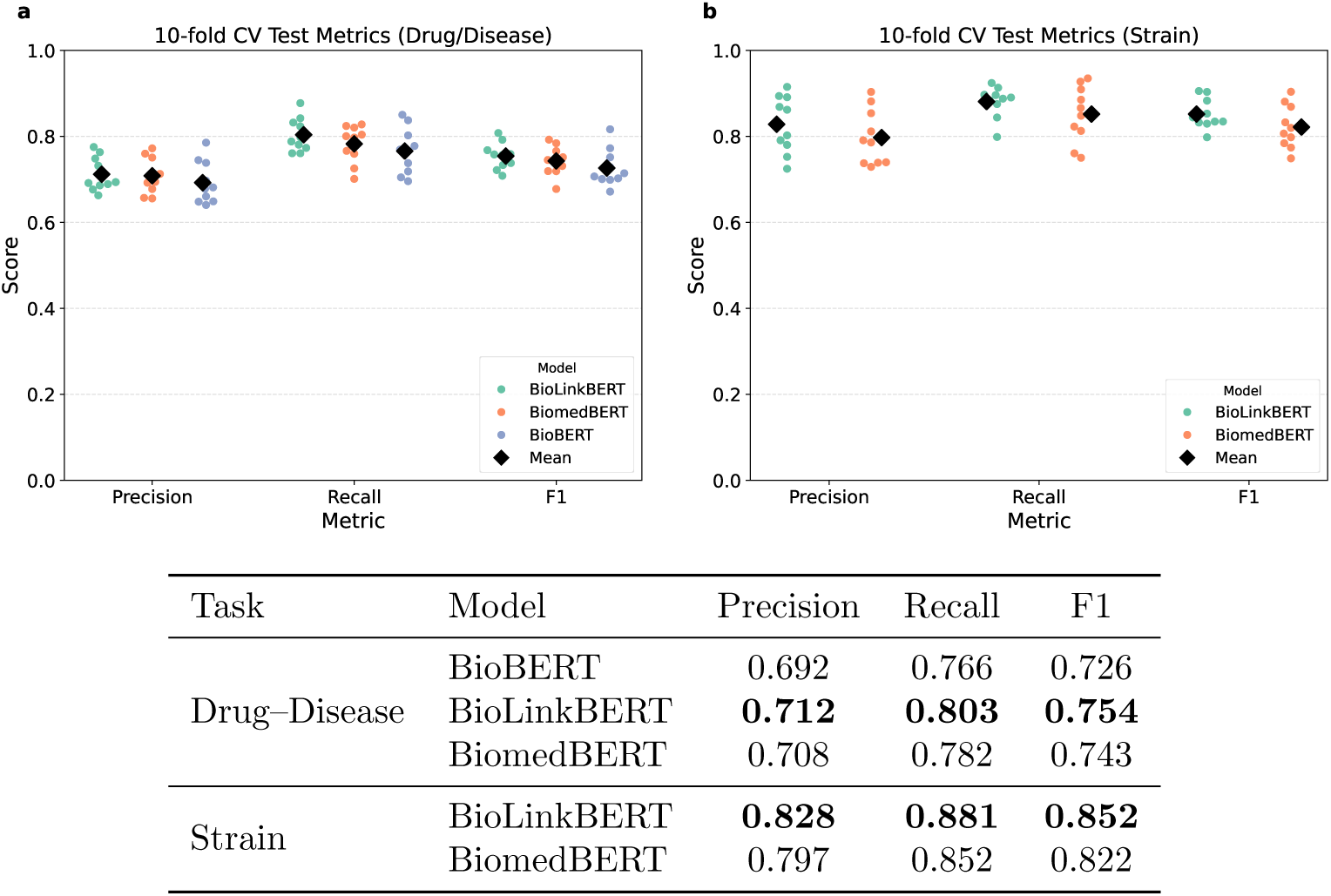
Performance of NER models using 10-fold cross-validation. Panels (a) and (b) show precision, recall, and F1 scores for drug–disease and strain annotation tasks, respectively. Points represent individual folds, and diamonds indicate mean performance. The table summarizes mean performance across folds for each model.

**Fig. C4:**
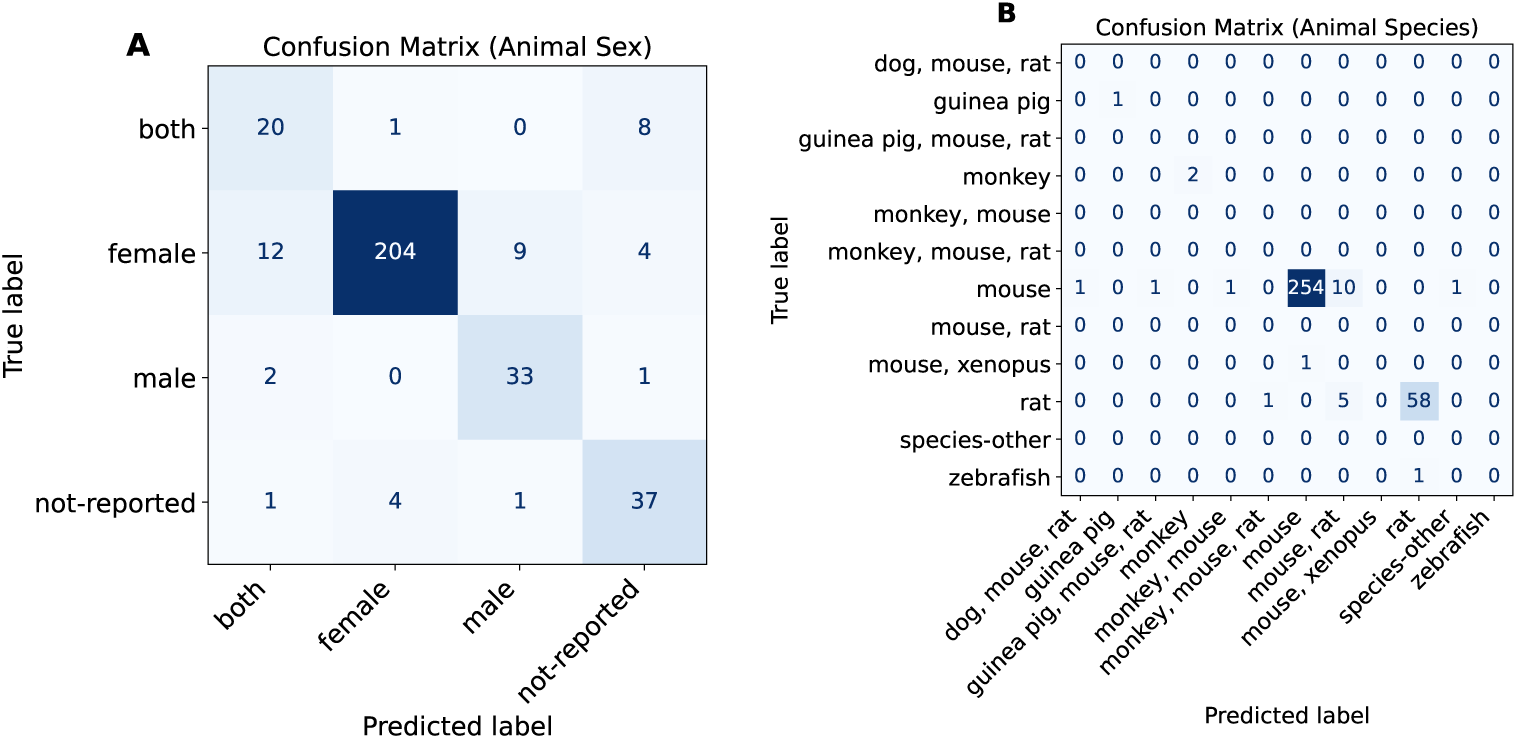
Confusion matrices for (A) animal sex and (B) species.

**Fig. C5:**
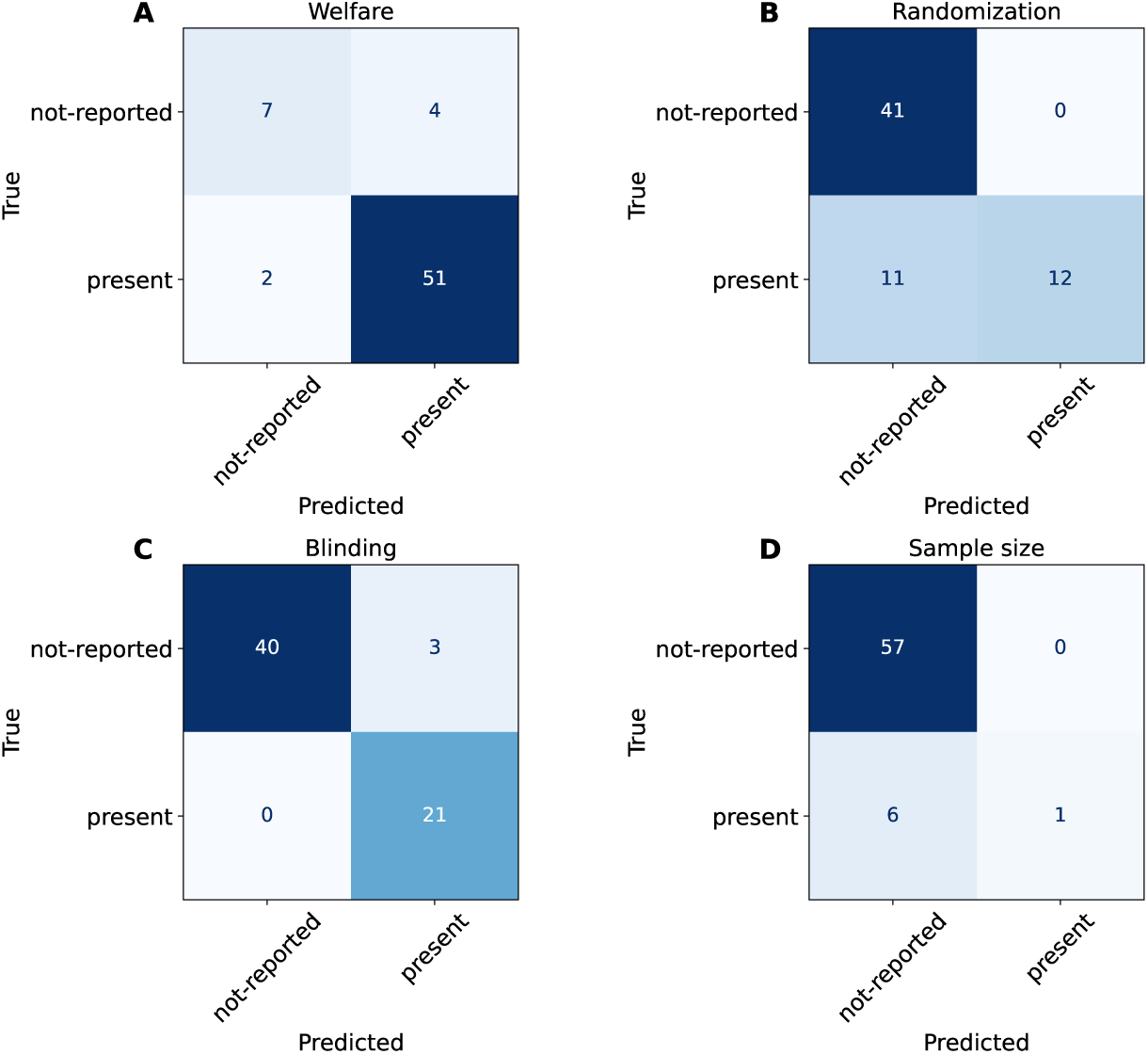
Confusion matrix for welfare statement and risk of bias items classification.

**Fig. E6:**
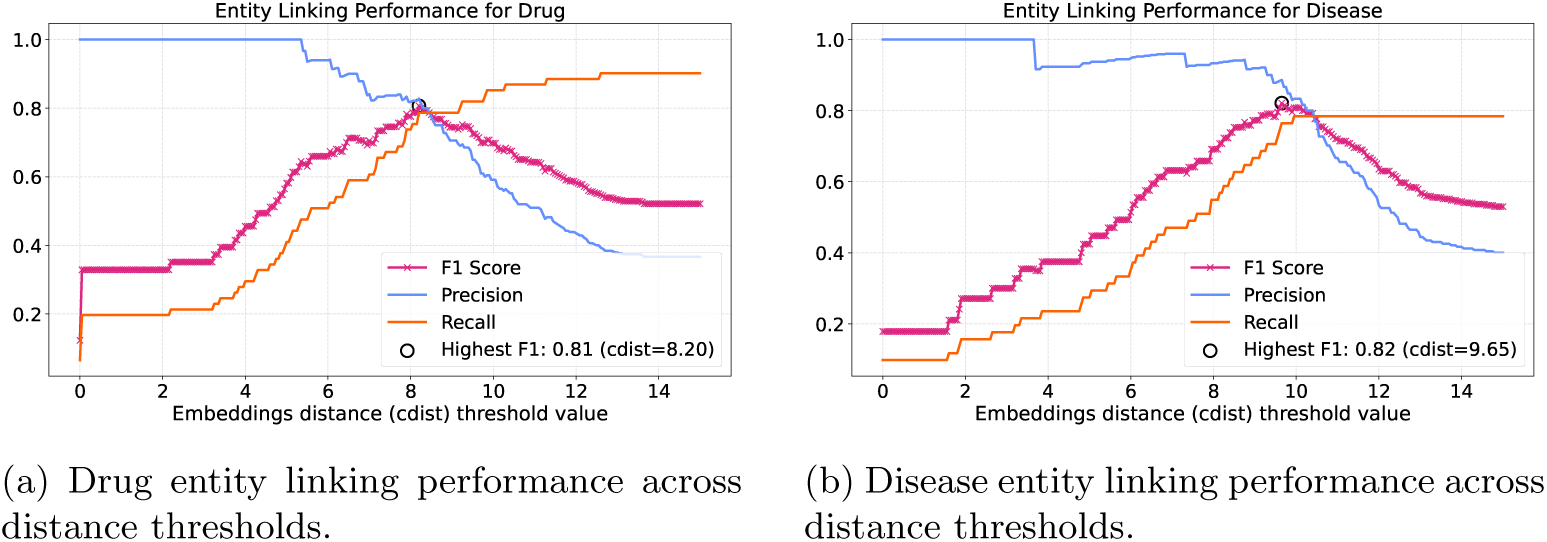
Entity linking performance as a function of the embedding distance (cdist) threshold.

**Fig. E7:**
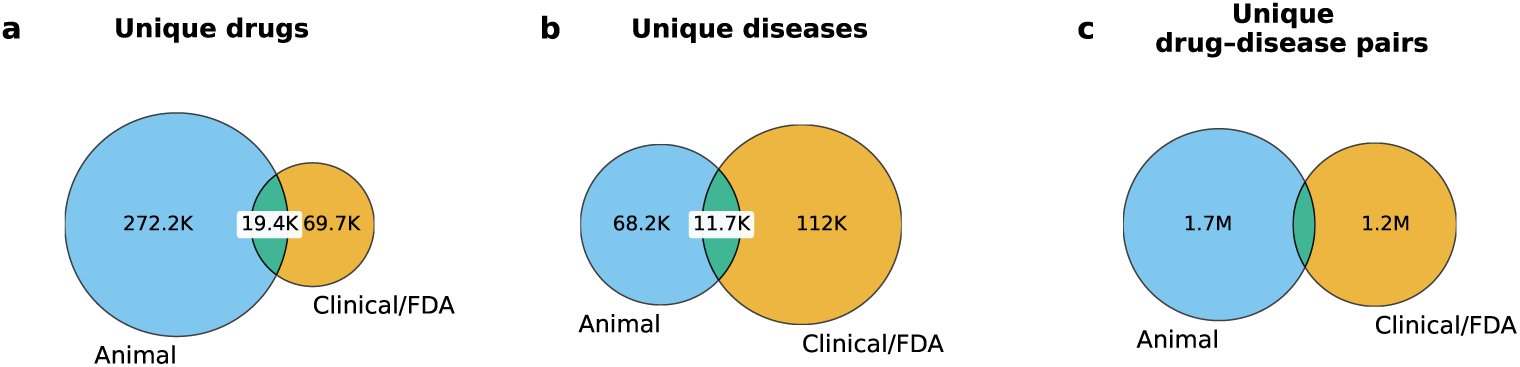
Venn diagrams illustrating the overlap between entities extracted from clinical/FDA data and those identified in preclinical (animal) studies. Panels show unique normalized (a) drugs, (b) diseases, and (c) drug–disease pairs. Numbers represent the counts of unique entities in each set and their intersections.

**Fig. F8:**
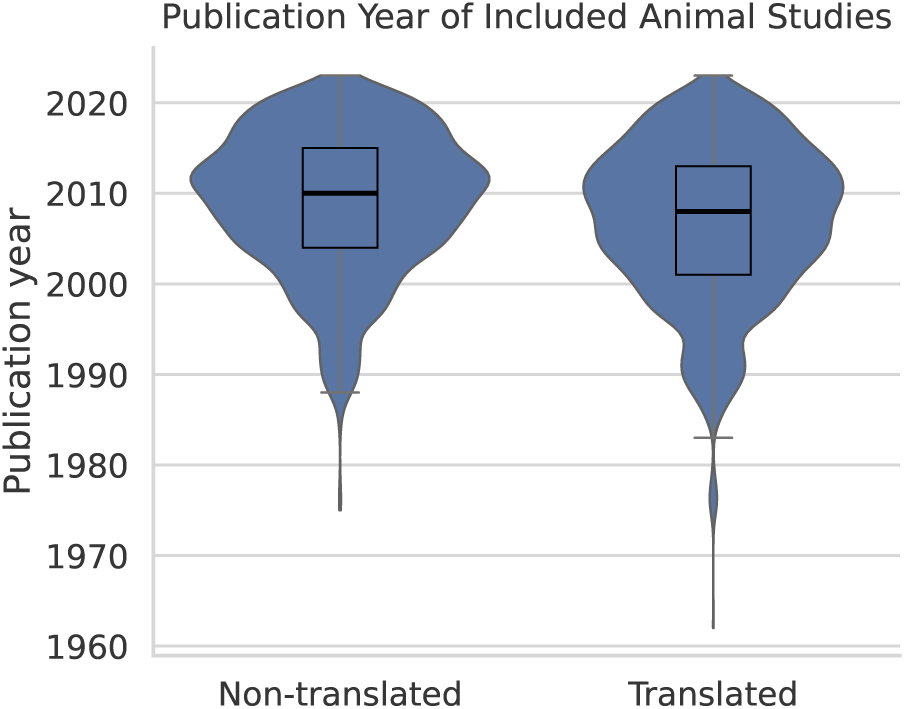
Distribution of publication years for included animal studies by outcome (translated vs. non-translated). Violin plots represent the density of publication years, with overlaid boxplots indicating the median and interquartile range.

**Fig. F9:**
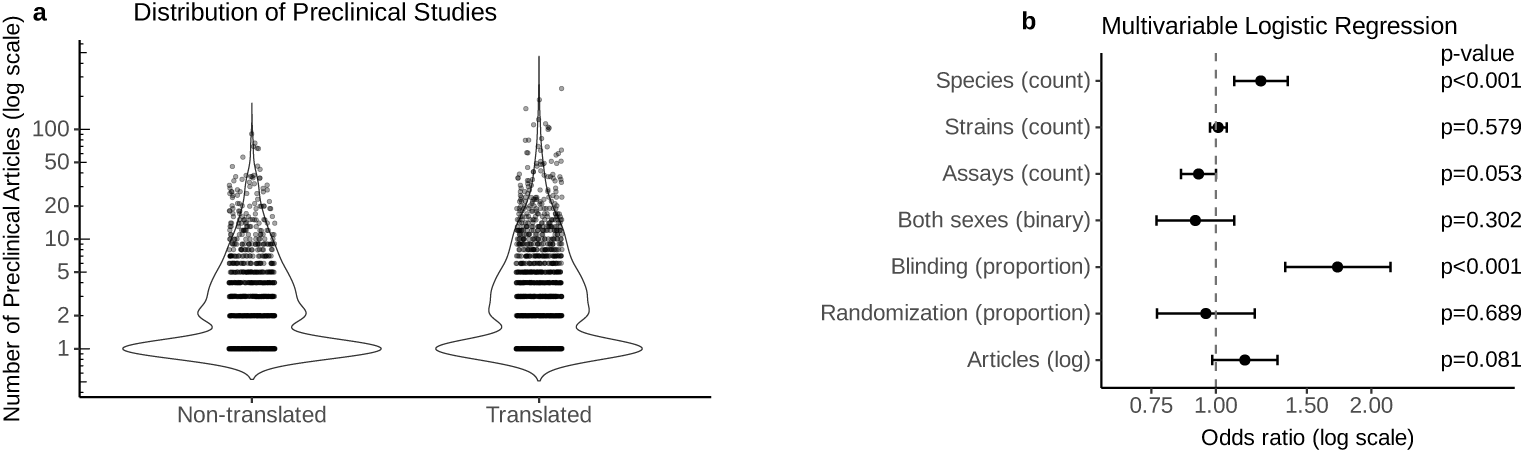
Preclinical evidence volume and its association with translation success. (a) Distribution of the number of supporting preclinical articles per drug–disease pair stratified by translation status (log scale). (b) Adjusted odds ratios from the multivariate logistic regression model including measures of external validity, internal validity, and evidence volume (log(articles)). Bars represent 95% confidence intervals.

**Fig. F10:**
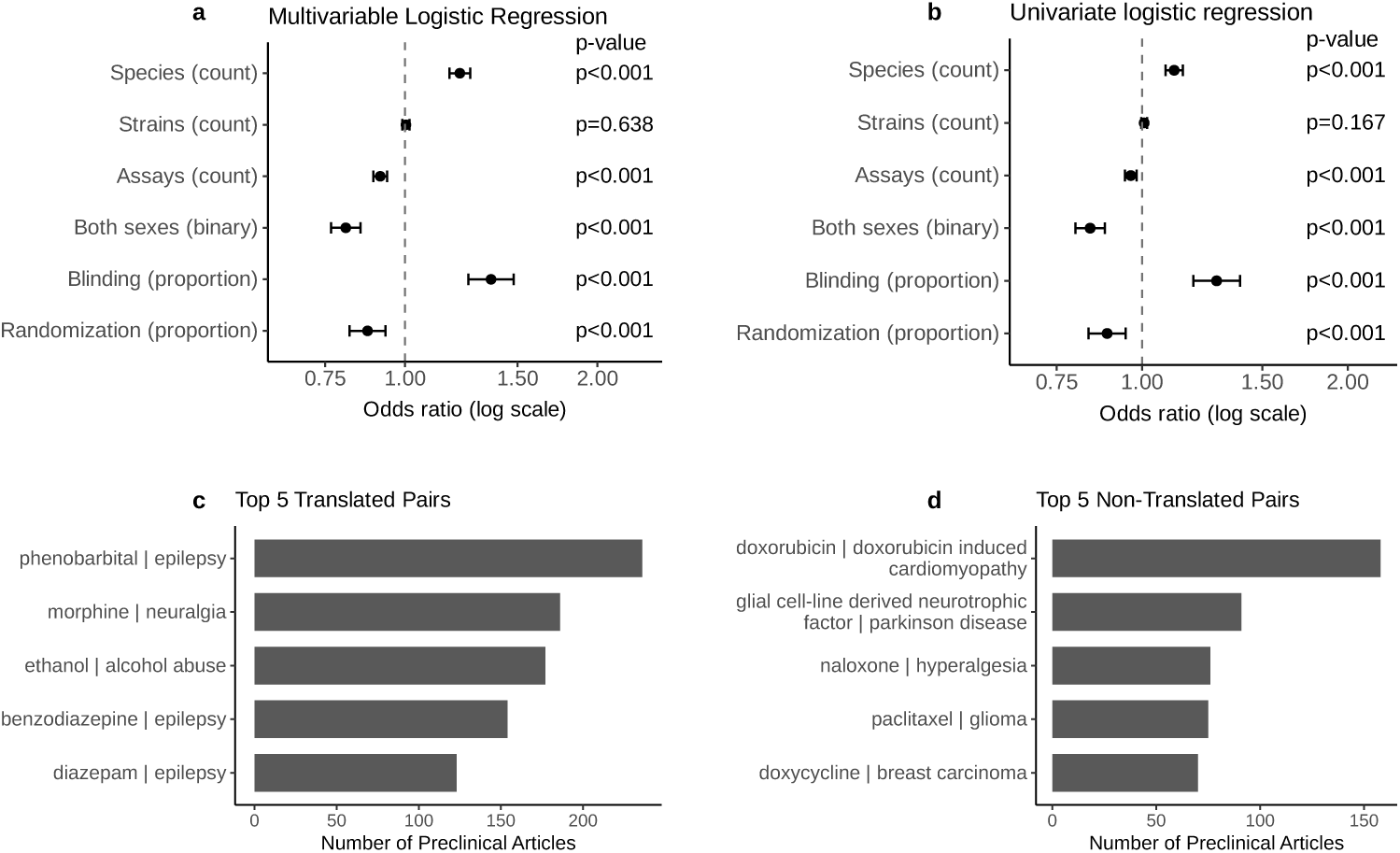
Associations between preclinical study characteristics and translation success for all drud-disease pairs. Adjusted odds ratios from multivariate (A) and univariate (B) logistic regression models. Panels (C) and (D) show the top 5 translated and non-translated drug–disease pairs ranked by the number of supporting preclinical articles. Bars indicate 95% confidence intervals in regression panels.

**Table B1:**
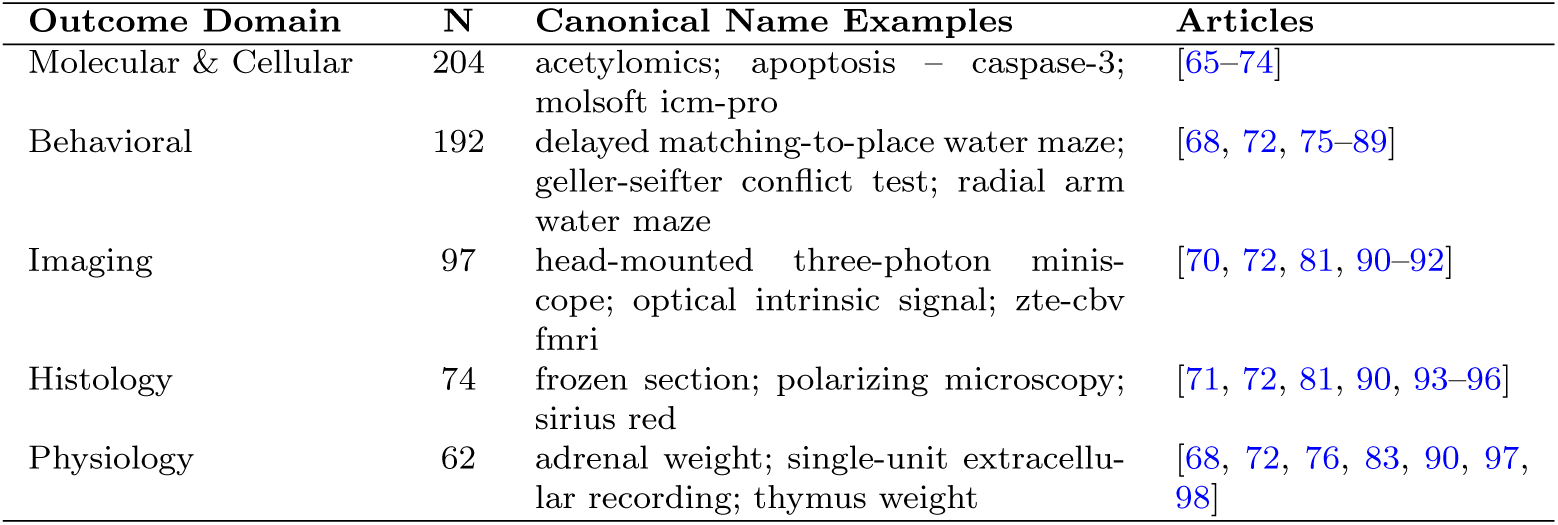
Summary of outcome domains, number of unique canonical names, and representative examples. Article references correspond to sources used during assay curation.

| Outcome Domain | N | Canonical Name Examples | Articles |
| --- | --- | --- | --- |
| Molecular & Cellular | 204 | acetylomics; apoptosis – caspase-3; molsoft icm-pro | [65–74] |
| Behavioral | 192 | delayed matching-to-place water maze; geller-seifter conflict test; radial arm water maze | [68, 72, 75–89] |
| Imaging | 97 | head-mounted three-photon miniscope; optical intrinsic signal; zte-cbv fmri | [70, 72, 81, 90–92] |
| Histology | 74 | frozen section; polarizing microscopy; sirius red | [71, 72, 81, 90, 93–96] |
| Physiology | 62 | adrenal weight; single-unit extracellular recording; thymus weight | [68, 72, 76, 83, 90, 97, 98] |

**Table C2:** Overview of extraction methods and performance metrics evaluated on the PreclinIE dataset. Methods include named entity recognition (NER) and rule-based regular expressions (regex). Metrics are reported as precision (P), recall (R), and F1-score (F1).

| Item | Approach | P | R | F1 |
| --- | --- | --- | --- | --- |
| Drug | NER → document-level aggregation | 0.78 | 0.85 | 0.81 |
| Disease | NER → document-level aggregation | 0.81 | 0.82 | 0.81 |
| Animal strain | NER → normalization | 0.83 | 0.88 | 0.85 |
| Animal species | Regex | 0.92 | 0.97 | 0.94 |
| Animal sex | Regex | 0.96 | 0.95 | 0.95 |
| Animal welfare | Regex | 0.98 | 0.96 | 0.97 |
| Randomization | Regex | 0.88 | 0.94 | 0.91 |
| Masking (blinding) | Regex | 0.92 | 0.93 | 0.93 |
| Sample size | Regex | 0.85 | 0.52 | 0.65 |

**Table C3:** Retention of target PMIDs across pipeline processing stages (n=463). Values show stage-specific and cumulative losses relative to the initial reference set.

| Stage | Included | Lost | Cum. lost | Remain. (%) |
| --- | --- | --- | --- | --- |
| Original PubMed query | 456 | 7 | 7 | 98.5 |
| Animal study type filter | 442 | 14 | 21 | 95.5 |
| NER for drug and disease identified | 427 | 15 | 36 | 92.2 |
| Full text retrieval | 337 | 90 | 126 | 72.8 |

**Table C4:**
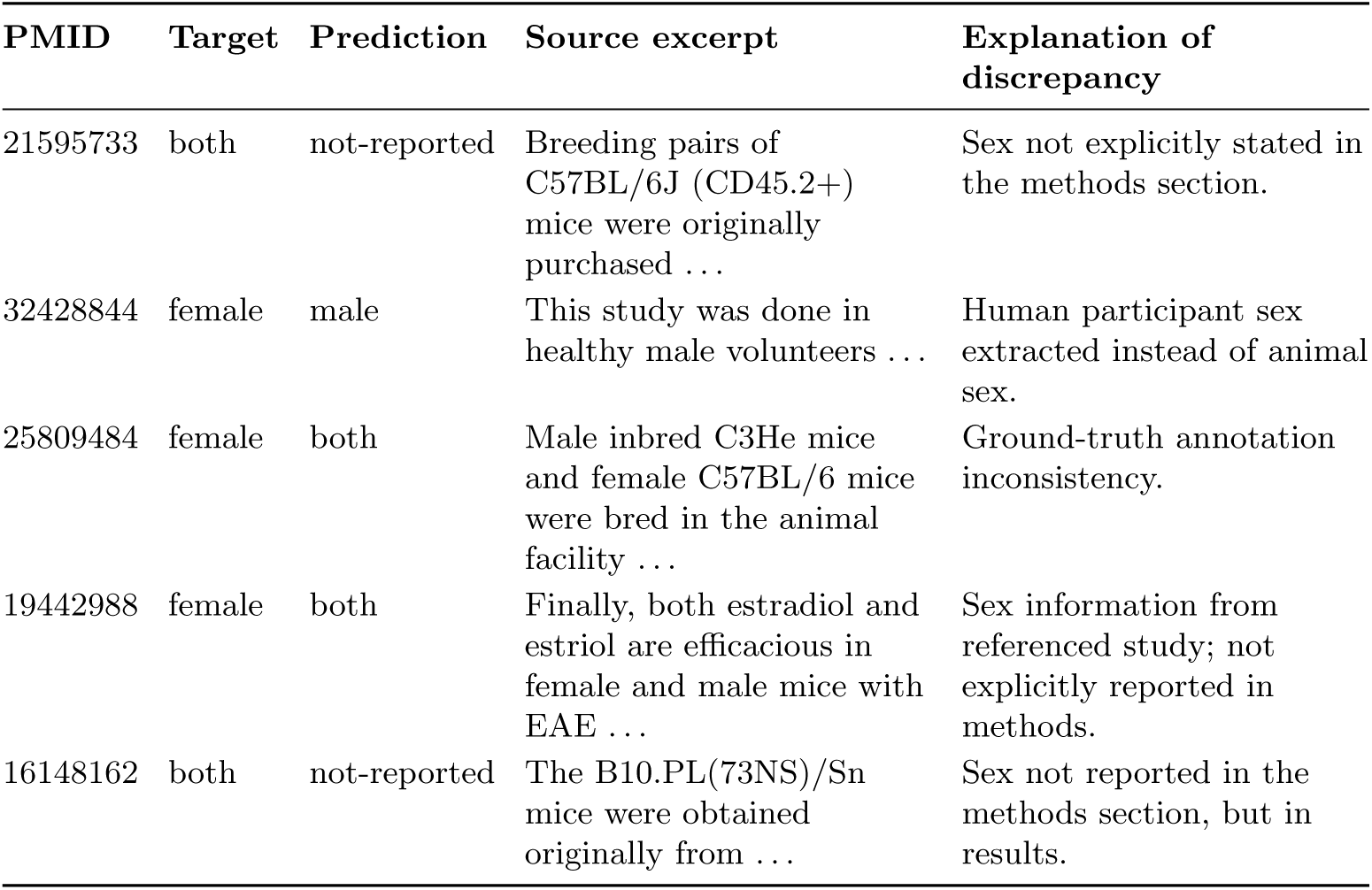
Manual review of five randomly sampled label–prediction discrepancies in the sex classification task.

**Table C5:** Manual review of five randomly sampled label–prediction discrepancies in the randomization reporting task.

| PMID | Target | Prediction | Source excerpt | Explanation of discrepancy |
| --- | --- | --- | --- | --- |
| 31844089 | present | not-reported | If multiple qualified patients were available for matching, random selection was employed ... | Information related to human study. Randomization not employed in the animal experiment. |
| 30858445 | present | not-reported | ...placed in the previous ‘forced’ runs (either left or right, randomly interchangeable between trials ... | Describes randomization of trial conditions rather than subject allocation. |
| 27078105 | present | not-reported | DMF treatment in randomized controlled MS trials was associated ... | Background sentence referring to human RCTs; randomization not employed in the animal experiment. |
| 27699272 | present | not-reported | No sentence containing the keyword found. | Target annotation source not clear. |
| 18540945 | present | not-reported | ...rats with protracted and severe EAE were randomized ... | Randomization mentioned in Results section rather than Methods. |

**Table D6:** Disease name extraction performance on the FDA indication dataset under exact and relaxed (substring-based) matching. Experiments were conducted using the vLLM framework with the DeepSeek-R1-Distill-Qwen-32B model.

| Match Type | Precision | Recall | F1 |
| --- | --- | --- | --- |
| Exact Match | 0.72 | 0.66 | 0.67 |
| Relaxed Match | 0.83 | 0.79 | 0.79 |

**Table D7:**
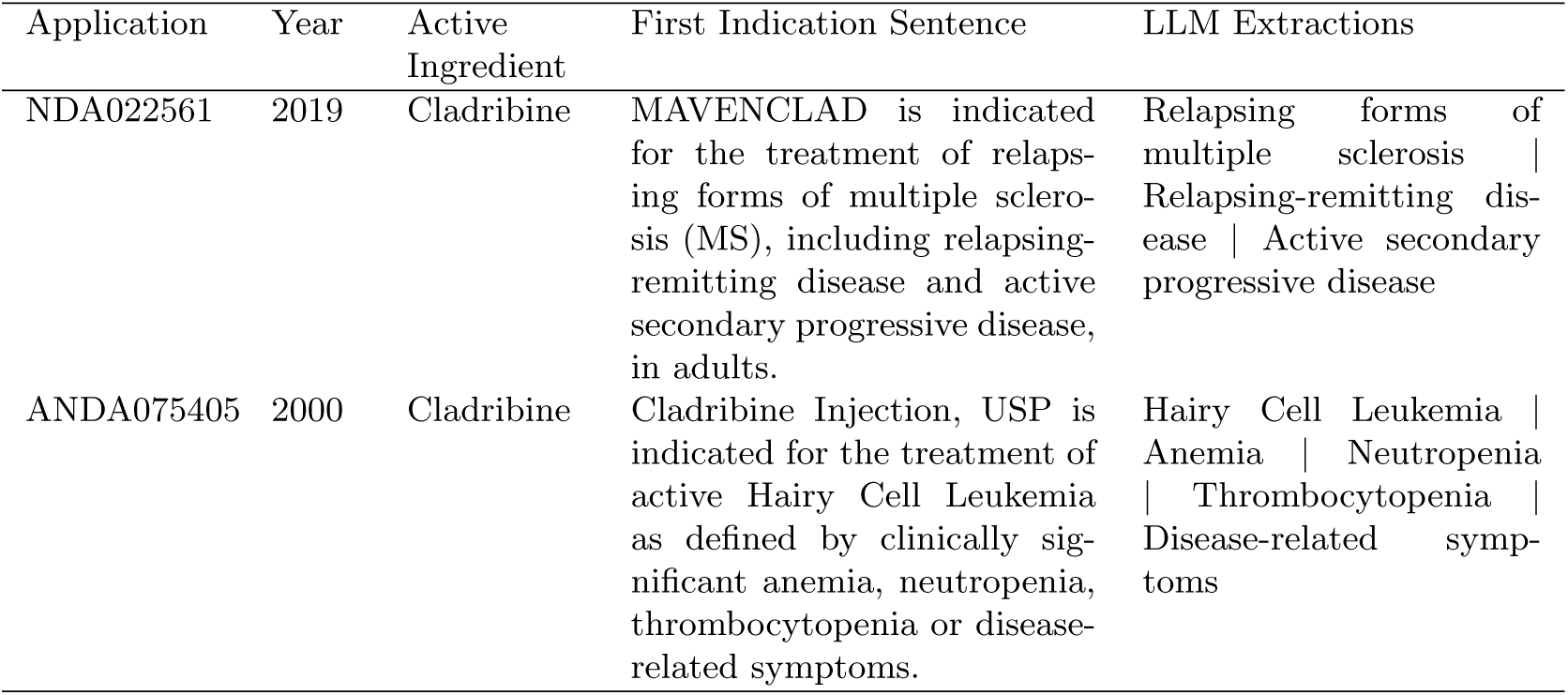
Example records obtained after merging Drugs@FDA application data with Drug Label indication text using application number as the linking key. The final column shows disease entities extracted from the indication sentence using a large language model (LLM).

**Table E8:** Overview of filtering rules applied to UMLS data.

| Filter | Field Name | Source Table | Target Values |
| --- | --- | --- | --- |
| Source Vocabulary | SAB | MRCONSO.RRF | ICD10, ICD10CM, ICD9CM, HPO, OMIM, SNOMEDCT_US, MSH, RXNORM, NDFRT, NDDF, MDR, DRUG-BANK, VANDF |
| Source Restriction Level | SRL | MRCONSO.RRF | 0, 9 |
| Is Preferred Name | ISPREF | MRCONSO.RRF | 1 |
| Term Type | TTY | MRCONSO.RRF | PT (Designated preferred name) |
| Drug-related Semantic Types | STY | MRSTY.RRF | Organic Chemical; Clinical Drug; Biologically Active Substance; Amino Acid, Peptide, or Protein; Enzyme; Immunologic Factor; Nucleic Acid, Nucleoside, or Nucleotide; Inorganic Chemical; Antibiotic; Biomedical or Dental Material; Hormone; Element, Ion, or Isotope; Drug Delivery Device; Vitamin; Chemical Viewed Structurally; Chemical |

**Table E9:**
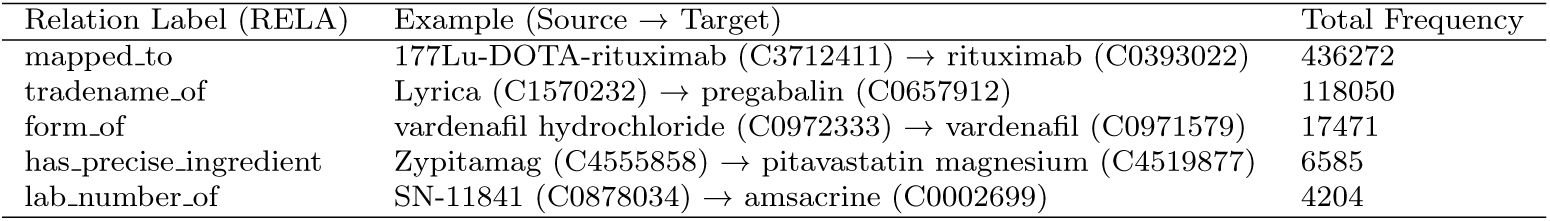
Examples of RN relations linking narrower drug terms to broader concepts, with total frequencies.

**Table E10:** Examples of normalized entities and their corresponding raw strings from the Named Entity Recognition (NER) step.

| Linked entity | NER raw string examples |
| --- | --- |
| **Drug entities (UMLS)** |  |
| MK-801<br>(C0813872) | “(+) mk-801”, “nmdar channel blocker mk-801”, “ss mk801” |
| Losartan<br>(C0126174) | “agtr1 inhibitor losartan”, “ang ii losartan”, “at1 blocker losartan” |
| Cannabinoid<br>(C0006864) | “type 1 cannabinoid”, “endogenous cannabinoid”, “cannabinoid receptor agon” |
| **Disease entities (MONDO)** |  |
| Brain ischemia<br>(MONDO:0005299) | “brain ischemia injury”, “brain ischemia-anoxia”, “ischemic cerebral disease” |
| Type 2 diabetes mellitus<br>(MONDO:0005148) | “2-diabetes”, “aging-related type 2 diabetes”, “adult onset dysglycemia” |
| Alzheimer disease<br>(MONDO:0004975) | “alzheimer disease (ad)”, “ad cognitive dysfunction”, “ad dementia” |

**Table E11:**
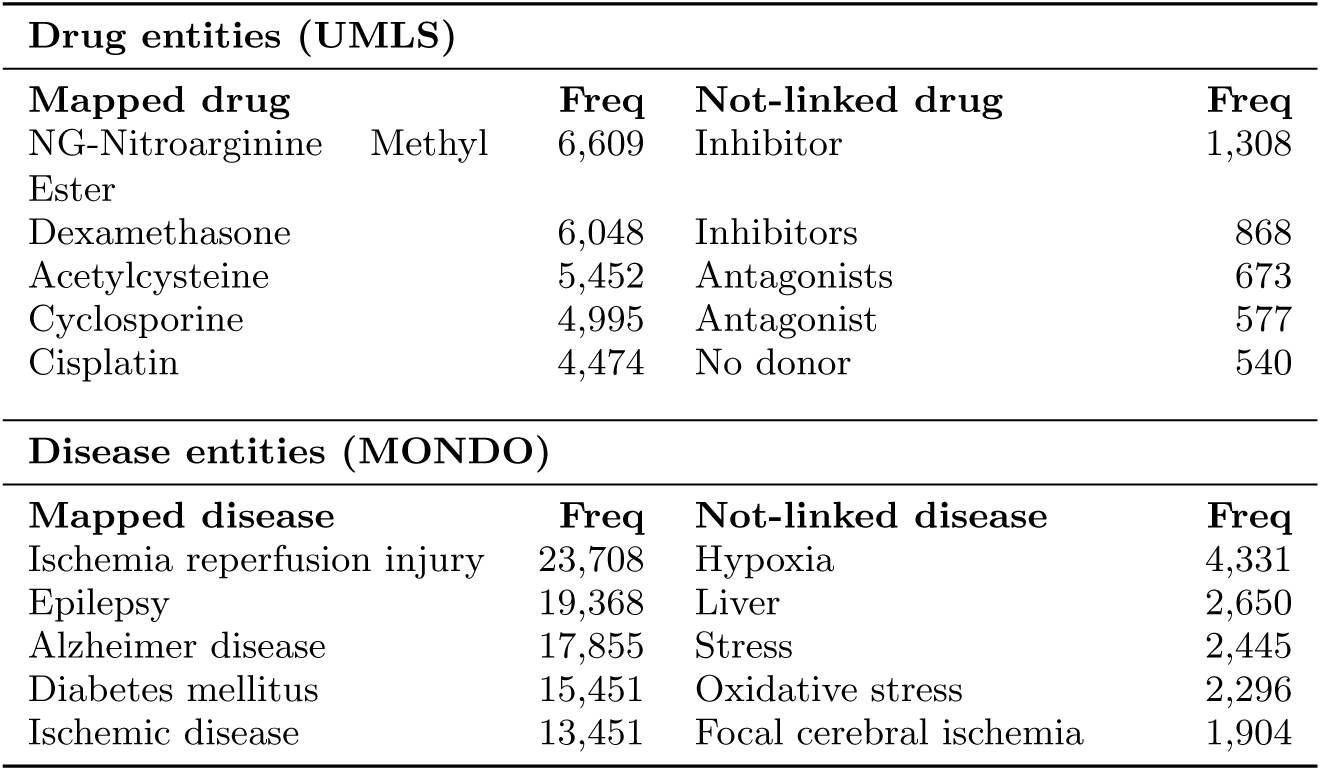
Top frequently mapped and not-linked drug and disease entities after normalization. Not-linked entities denote mentions for which no ontology concept met the similarity threshold.

**Table F12:**
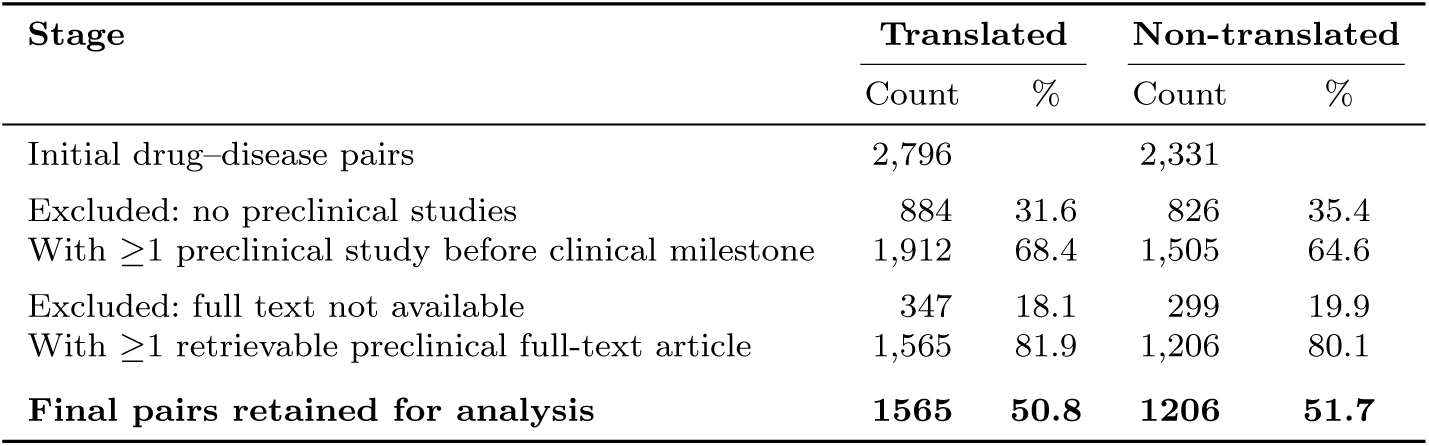
Preclinical retrieval funnel for translated and non-translated drug–disease pairs. Counts and percentages show progressive exclusions and retention based on the availability of preclinical evidence and retrievable full-text articles.

| Stage | Translated |  | Non-translated |  |
| --- | --- | --- | --- | --- |
|  | Count | % | Count | % |
| Initial drug-disease pairs | 2,796 |  | 2,331 |  |
| Excluded: no preclinical studies | 884 | 31.6 | 826 | 35.4 |
| With $\geq 1$ preclinical study before clinical milestone | 1,912 | 68.4 | 1,505 | 64.6 |
| Excluded: full text not available | 347 | 18.1 | 299 | 19.9 |
| With $\geq 1$ retrievable preclinical full-text article | 1,565 | 81.9 | 1,206 | 80.1 |
| **Final pairs retained for analysis** | **1565** | **50.8** | **1206** | **51.7** |

**Table F13:** Illustrative example of study-level records and their aggregation into a single pair-level observation for the drug–disease pair sildenafil *<>* traumatic brain injury (TBI).

| PMID | Species | Sex | Strain | Assay | Country |
| --- | --- | --- | --- | --- | --- |
| *Study-level records* |  |  |  |  |  |
| 30829389 | rat | female | Wistar | Histology | Turkey |
| 24662845 | rat | male | Sprague-Dawley | Behavioral;<br>Molecular;<br>Physiology | USA |
| *Aggregated across supporting studies (pair-level)* |  |  |  |  |  |
| sildenafil <> TBI | 1 unique | both | 2 unique | 4 unique | 2 unique |

**Table F14:**
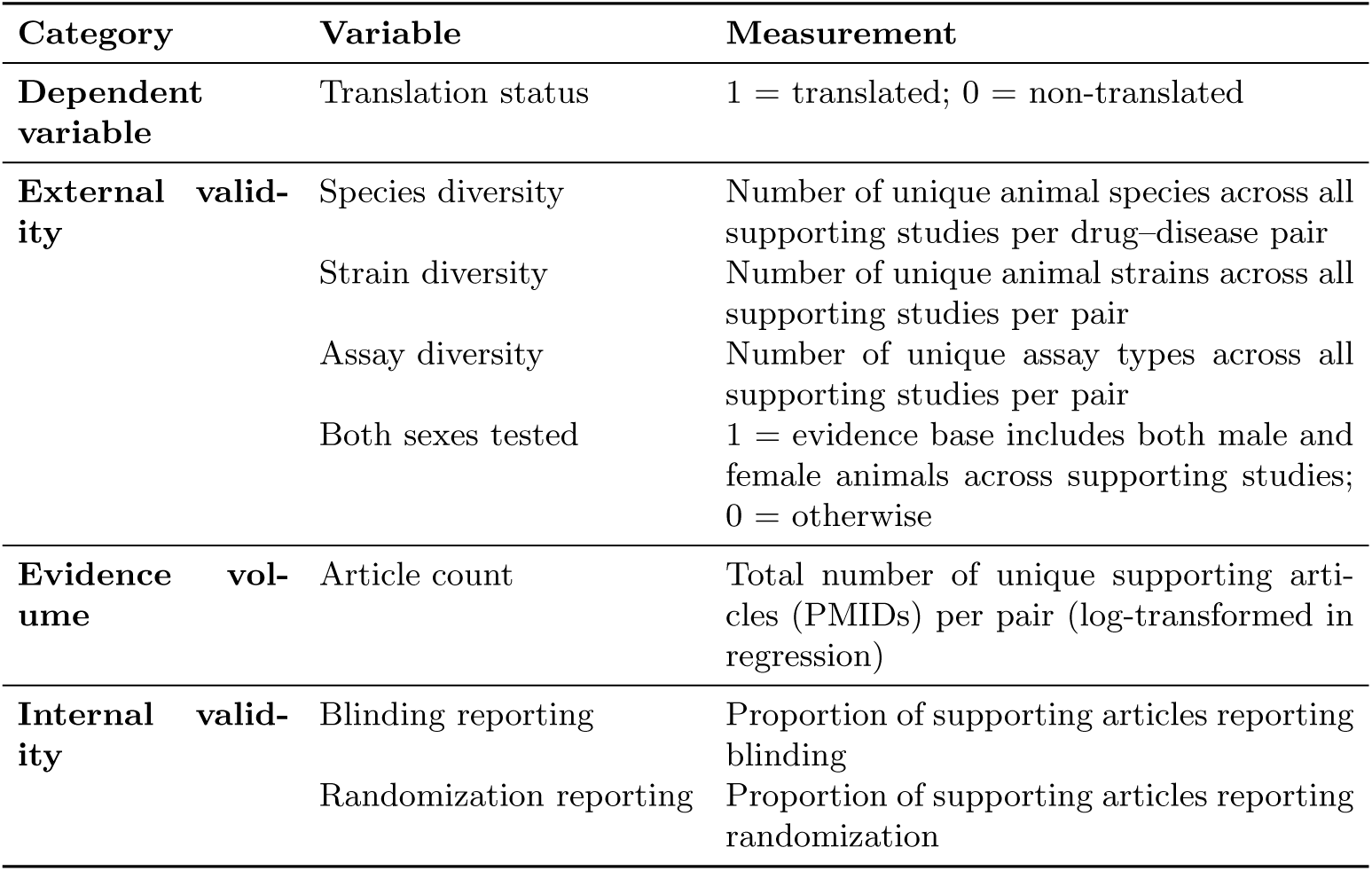
Variables used in the logistic regression analysis.

| Category | Variable | Measurement |
| --- | --- | --- |
| **Dependent variable** | Translation status | 1 = translated; 0 = non-translated |
| **External validity** | Species diversity | Number of unique animal species across all supporting studies per drug–disease pair |
|  | Strain diversity | Number of unique animal strains across all supporting studies per pair |
|  | Assay diversity | Number of unique assay types across all supporting studies per pair |
|  | Both sexes tested | 1 = evidence base includes both male and female animals across supporting studies; 0 = otherwise |
| **Evidence volume** | Article count | Total number of unique supporting articles (PMIDs) per pair (log-transformed in regression) |
| **Internal validity** | Blinding reporting | Proportion of supporting articles reporting blinding |
|  | Randomization reporting | Proportion of supporting articles reporting randomization |

**Table F15:** Univariate and multivariable logistic regression analyses examining associations between preclinical study characteristics and translation of drug–disease pairs. Odds ratios greater than 1 indicate higher odds of translation.

| Predictor | Univariate model |  | Multivariable model |  |
| --- | --- | --- | --- | --- |
| | OR (95% CI) | $p$ | AOR (95% CI) | $p$ |
| Species tested (count) | 1.28 (1.18–1.40) | <0.001 | 1.26 (1.13–1.42) | <0.001 |
| Strains tested (count) | 1.06 (1.03–1.08) | <0.001 | 1.03 (1.00–1.06) | 0.096 |
| Assays tested (count) | 1.08 (1.03–1.14) | 0.004 | 0.96 (0.89–1.02) | 0.199 |
| Tested in both sexes (binary) | 1.13 (0.97–1.32) | 0.107 | 0.93 (0.79–1.11) | 0.425 |
| Blinding reported (proportion) | 1.61 (1.29–2.01) | <0.001 | 1.68 (1.33–2.12) | <0.001 |
| Randomization reported (proportion) | 1.04 (0.84–1.28) | 0.711 | 0.95 (0.76–1.18) | 0.644 |

**Table F16:**
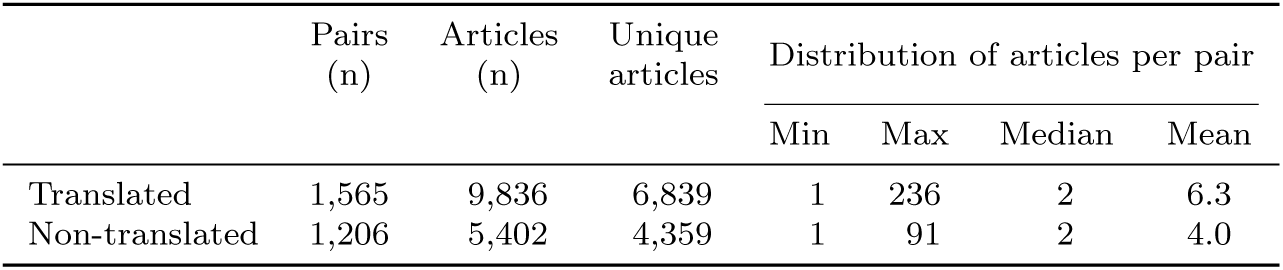
Summary statistics of supporting preclinical articles per drug–disease pair according to translation status.

## Footnotes

1 Detailed search queries: https://github.com/Ineichen-Group/Preclinical_Pipeline/tree/main/01_pubmed_query_neuro/data/pubmed_queries.

2 Database snapshot, 1 December 2025. Available at https://aact.ctti-clinicaltrials.org/downloads/snapshots?type=pgdump&year=2025.

3 Accessed 11 November 2025 from https://open.fda.gov/

4 https://huggingface.co/cambridgeltl/SapBERT-from-PubMedBERT-fulltext

5 Included diseases: multiple sclerosis, Alzheimer’s, Parkinson’s, and Huntington’s diseases; motor neuron disease (including amyotrophic lateral sclerosis); neuropathic pain; prion disease (including creutzfeldt); spinal cord injury; traumatic brain injury; subarachnoid hemorrhage; stroke; schizophrenia; depression; addiction; autism spectrum disorder; brain tumors (glioblastoma, glioma, meningioma); and epilepsy.

6 https://github.com/Ineichen-Group/HERMES-MS/blob/main/Files-backup/HERMES_INCLUDED.csv

7 https://github.com/Ineichen-Group/HERMES-MS/blob/main/mined_rob_with_addpaper.xlsx

8 https://www.nlm.nih.gov/research/umls/rxnorm/sourcereleasedocs/drugbank.html

9 /cambridgeltl/SapBERT-from-PubMedBERT-fulltext

